# EARLY AND MID-GESTATION ZIKA VIRUS (ZIKV) INFECTION IN THE OLIVE BABOON (*PAPIO ANUBIS*) LEADS TO FETAL CNS PATHOLOGY BY TERM GESTATION

**DOI:** 10.1101/2022.02.23.481575

**Authors:** Sunam Gurung, Darlene Reuter, Abby Norris, Molly Dubois, Marta Maxted, Krista Singleton, Marisol Castillo-Castrejon, James F. Papin, Dean A. Myers

## Abstract

Zika virus (ZIKV) infection in pregnancy can produce catastrophic teratogenic damage to the developing fetus including microcephaly and congenital Zika syndrome (CZS). We previously described fetal CNS pathology occurring by three weeks post-ZIKV infection in Olive baboons at mid-gestation, including neuroinflammation, loss of radial glia (RG), RG fibers, neuroprogenitor cells (NPCs) resulting in disrupted NPC migration. In the present study, we explored fetal brain pathologies at term gestation resulting from ZIKV infection during either first or second trimester in the Olive baboon. In all dams, viremia resolved after 7 days post infection (dpi). One first trimester infected dam aborted at 5 dpi. All dams developed IgM and IgG response to ZIKV with ZIKV IgG detected in fetal serum. Substantial placental pathology and inflammation was observed including disruption of syncytiotrophoblast layers, delayed villous maturation, partially or fully thrombosed vessels, calcium mineralization and fibrin deposits. In the uterus, ZIKV was detected in ¾ first trimester but not in second trimester infected dams. While ZIKV was not detected in any fetal tissue at term, all fetuses exhibited varying degrees of neuropathology. Fetal brains from ZIKV infected dams exhibited a range of gross brain pathologies including irregularities of the major gyri and sulci of the cerebral cortex and cerebellar pathology. Frontal cortices of ZIKV fetuses showed a general disorganization of the six-layered cortex with degree of disorganization varying among the fetuses from the two groups. Frontal cortices from first but not second trimester infections exhibited increased microglia and astrocyte numbers (white matter) in both trimester infections. In the cerebellum, increased microglia were observed in both first and second trimester infected fetuses and decreased oligodendrocyte precursor cell populations in the cerebellar white matter in first trimester infections. In general, our observations are in accordance with those described in human ZIKV infected fetuses.

## INTRODUCTION

The epidemic potential and transmission of emerging and re-emerging flaviviral diseases such as Zika virus (ZIKV) pose a major threat to the global public health and economy. Besides ZIKV, other flaviviruses of great public health concern belonging to the *Flaviviridae* family include dengue (DENV), West Nile (WNV), yellow fever (YFV), and Japanese encephalitis virus (JEV) [1, 2]. While ZIKV infections have declined since its peak in ∼2017, where it had spread to 84 countries with over 800,000 cases reported in the Americas alone (PAHO), it remains a global threat due to continued presence in tropical American regions, lack of a vaccine, and continued expansion of its primary mosquito vector, *Aedes aegypti*. Although ZIKV infection is typically mild or asymptomatic, with symptoms including minor fever, rash and conjunctivitis, ZIKV can exert major pathologies on pregnancy including spontaneous abortion, intrauterine fetal death, fetal growth restriction, microcephaly and associated neurological damage. Collectively these effects on the fetus are termed congenital zika syndrome (CZS) [3, 4]. Newborns of women infected with ZIKV during pregnancy have a 5-14% risk of developing CZS [5]. It was estimated that 10% of babies born to infected mothers in Brazil in 2015 had some birth defect [6].

Studies show poor developmental prognosis for children with CZS with nearly 100% presenting with severe developmental injuries by 30 months age [7]. Children born from ZIKV infected mothers have been reported to exhibit below average neurodevelopmental assessment (cognitive, language, motor) between 7 and 32 months of age, and some developed autism spectrum disorder in the second year of life [8]. These neurological, musculoskeletal, cognitive and developmental complications have also been reported in infants born without clinical abnormalities at birth [9, 10] suggesting continued detrimental developmental consequences in response to prenatal ZIKV exposure. In the United States, 9% of children born to ZIKV infected mothers were reported to have at least one neurodevelopmental abnormality before they reached 2 years of age [11].

CNS pathologies resulting from ZIKV infection in pregnancy include structural (thin cerebral cortices with subcortical calcifications, ventriculomegaly, cerebellar hypoplasia and atrophy, lissencephaly, pachygyria/agyria, hydrocephaly, and various cortical development errors), ocular (cataract, glaucoma, optic nerve abnormalities, focal pigment mottling, intraocular calcifications, chorioretinal macular atrophy/scarring), congenital contractures (arthrogryposis), hypotonia and clinical signs such as craniofacial malformations, swallowing disorders, auditory and sensorineural defects [12]. Postnatal development of microcephaly has also been observed in infants born with normal head circumference [13].

The mechanisms driving the severe neurological damage to the developing human fetus from a ZIKV infected mother have not been fully elucidated. Therefore, animal models serve as essential surrogates to study the pathogenesis of ZIKV during pregnancy. Mouse models of ZIKV infection have been widely leveraged to understand pathogenesis of ZIKV induced neural damage. For example, Yuan et al, reported that a single mutation (S139N) in the ZIKV PrM protein acquired in the French Polynesian ZIKV isolate resulted in more efficient ZIKV infection of neural progenitor cells (NPCs) in the fetal mouse brain compared to the ancestral Asian ZIKV isolate leading to significant disturbance of neurogenesis and microcephaly [14]. Studies in ZIKV infected mice have shown downregulation of genes involved in cell proliferation, differentiation, neurogenesis, migration as well as genes involved in DNA repair and replication [15, 16]. Similar downregulation of genes involved in neurogenesis and survival were also seen in response to ZIKV in human NPCs, neurospheres and organoids [17]. In fetal mice, ZIKV has been shown to target glial and neuronal population including NPCs, neurons, oligodendrocyte progenitor cells (OPCs), astrocytes and microglia. Neuronal cell death, microglial hyperplasia, gliosis with reactive astrocytes, and myelination delay have been reported in postnatal mice [18, 19]. The same neurological pathogenesis has been reported in ZIKV infected human microcephalic fetuses [20, 21]. Despite all of the significant insights about ZIKV induced neurological pathogenesis gained from the mouse models, it poses significant challenges to model human pregnancy due to differences in gestation, placentation and fetal brain development. Wild-type mice are also resistant to ZIKV infection; hence, most studies in mice have relied on mice deficient in interferon signaling or using direct routes of inoculation (e.g., intrauterine, vaginal) to achieve ZIKV pathogenesis thus questioning the translational applicability of the mouse studies [22–26].

Being natural animal reservoirs for Zika and ZIKA-related flaviviruses, non-human primates (NHPs) are excellent translational models for studying ZIKV pathogenesis. Zika virus infection has been extensively studied in baboons [27–30], macaques (rhesus [31–37], pigtail [38, 39], cynomolgus [40, 41]), and marmosets (*Callithrix jacchus;* [42]). Similar to humans, intrauterine fetal death and/or miscarriage has been reported as a common (26%) outcome following ZIKV infection in macaques and baboons [43]. While microcephaly has not been reported in macaques infected with ZIKV, a range of fetal or infant neuropathological outcomes have been described from none [31, 44] to notable [35, 38], including CNS lesions (microcalcifications, hemorrhage, vasculitis, necrosis), gliosis (microglia, astrocytes), white matter injury, calcification, loss of NPCs, reduced brain size (but not microcephaly) and agyria, with the latter being rare [35, 38, 39, 45]. Similarly, a range of placental pathology has been noted in macaques from negligible [38, 44] to severe placental damage and placental insufficiency [33, 35]. Hypoxia and inflammation in the developing fetus due to placental/vascular insufficiency may contribute to fetal brain damage and CZS [35]. ZIKV infection early in pregnancy, compared to later gestation, is associated with more severe fetal brain damage and CZS in macaques consistent with human data [46]. Cumulatively, studies in rhesus macaques have shown a high (100%) rate of vertical transfer of ZIKV with a diverse range of fetal/infant neuropathology, unlike humans where the rate of vertical transfer is estimated between 10-20%.

We developed an olive baboon (*Papio anubis*) model to study ZIKV pathogenesis, The olive baboon is similar to humans in terms of size, genetics, placentation, gestation, brain development and immune repertoire making the baboon an excellent translational NHP model to study ZIKV infection and for vaccine and therapeutics development [47–49]. The baboon is permissive to flavivirus infection and replication, including ZIKV, and produces a virus-specific immune response [49, 50]. Our previous study focused on the early events of infection during the three weeks post-subcutaneous ZIKV infection of pregnant dams during mid-gestation to decipher the timing of vertical transfer and early events of fetal CNS targeting [28]. Prior studies in macaques typically examined fetal (and placental) pathologies at term gestation or early post-delivery. We observed vertical transfer in ½ of the infected dams. In one fetus at 3 weeks post-maternal infection, we observed extensive neuropathology including a significant loss in radial glia (RG) and their fibers, decreased NPCs, astrogliosis and increased reactive microglia as well as decreased oligodendrocyte precursor cells (OPCs). We also noted disorganization of neurons in the cortical plate, consistent with the loss of RG fibers, which serve as scaffolding for newly formed neurons and NPCs during migration to the cortical plate forming the characteristic six-layered cortex.

To our knowledge, we were the first to report the loss of RG fibers due to ZIKV infection in primates, which had been previously seen in microcephalic human fetuses and in mouse models [28]. The present study extended these findings by examining fetal neurological outcome at term gestation following ZIKV infection during late first trimester or at mid-gestation in the olive baboon.

Herein, we describe infection of four timed-pregnant olive baboons at early-gestation and three at mid-gestion with a contemporary Puerto Rican strain of ZIKV (PRVABC59, 2015). We report extensive fetal brain pathology from both early and mid-gestation infected dams, one fetal abortion in the early gestation group, viral persistence of ZIKV RNA in the uterus of early gestation infected dams and in various lymph nodes in both the early and mid-gestation infected dams at the time of termination of the study.

## RESULTS

### Symptomology and pregnancy outcome

In the early-gestation (E) cohort, Dam 1E exhibited slight erythema in the inguinal, axillary and injection site areas on 7 dpi which progressed to mild maculopapular rash in axillary and moderate to severe in inguinal areas at 14 and 21 dpi while resolving to mild in both areas by day 40 dpi. However, by 80 dpi, this animal exhibited moderate to severe maculopapular rash in the axillary area. Dam 2E exhibited only slight erythema at 14 dpi in the inguinal area that resolved by later time points. Dam 3E exhibited slight erythema in the inguinal area at 80 dpi. Dam 4E did not exhibit clinical signs before aborting at 5 dpi.

In the mid-gestation (M) cohort, Dam 1M exhibited clinical signs throughout pregnancy, mainly slight erythema in axillary and inguinal areas, localized redness in upper abdomen and chest and significant conjunctivitis on 4 to 42 dpi, with resolution of conjunctivitis by 42 dpi. Dam 2M exhibited slight axillary and inguinal erythema and mild conjunctivitis which were resolved by 40 dpi. Dam 3M exhibited slight axillary and inguinal erythema and slight to mild conjunctivitis at 7dpi which resolved by 21 dpi. None of the dams exhibited body temperatures (obtained under ketamine sedation) greater than 1°C above day 0 over the course of the study.

### Maternal ZIKV Viral loads

In the early-gestation cohort, viremia (ZIKV RNA) initiated at 4 dpi in Dams 1E, 2E, 3E and 4E. Dams 1E and 2E were also viremic at 7 dpi but not at later time points. ZIKV RNA was detected in saliva from Dams 1E and 3E at 4 dpi (Dam 1E), 4, 7 dpi (Dam 2E) and 7 dpi (Dam 3E) (**Table 1**). We did not detect ZIKV RNA in vaginal swabs obtained at any time point for Dams 1E and 2E. ZIKV RNA was detected in vaginal swabs from Dam 3E at 4, 7, 14 and 21 dpi. Dam 4E aborted 5 dpi, therefore, the final samples for this dam were collected during necropsy on day 5 post infection.

**Table 1.** ZIKV RNA in select maternal, fetal tissues (copies per mg tissue or per ml of fluid)

| Fluids/Tissues | Dam 1E (Early) | Dam 2E (Early) | Dam 3E (Early) | Dam 4E (Early) | Dam 1M (Mid) | Dam 2M (Mid) | Dam 3M (Mid) | Control (n=3) |
| --- | --- | --- | --- | --- | --- | --- | --- | --- |
| Gestation day start/end | 56-172 dG | 58-169 dG | 55-169 dG | 55-59 dG | 92-167 dG | 94-169 dG | 97-167 dG | 169, 171, 166 dG |
| **Maternal Samples** |  |  |  |  |  |  |  |  |
| Whole Blood | 10<sup>3</sup> (D4) | 10<sup>4</sup> (D4) | 10<sup>4</sup> (D4) | 10<sup>4</sup> (D4) | 10<sup>3</sup> (D7) | <LD | 10<sup>5</sup> (D4) | - |
| Saliva | 10<sup>3</sup> (D4) | 10<sup>4</sup> (D4) | 10<sup>3</sup> (D7) | <LD | <LD | 10<sup>3</sup> (D7) | 10<sup>5</sup> (D7) | - |
| Vaginal Swab | <LD | <LD | 10<sup>5</sup> (D14) | <LD | <LD | <LD | <LD | - |
| Uterus | 10<sup>4</sup> | <LD | 10<sup>6</sup> | 10<sup>6</sup> | <LD | <LD | <LD | - |
| Lymph node | 10<sup>3</sup> | <LD | <LD | 10<sup>5</sup> | 10<sup>4</sup> | 10<sup>3</sup> | 10<sup>4</sup> | - |

| Fetal Samples |  |  |  |  |  |  |  |  |
| --- | --- | --- | --- | --- | --- | --- | --- | --- |
| Fetal weight | 1.06 kg | 0.665 kg | 1.06 kg | N/A | 0.885 kg | 0.965 kg | 0.780 kg | 0.80, 1.05, 0.8 kg |
| Viability | V | V | V | Aborted | V | V | V | V |
| Sex | F | F | M | - | M | F | F | M, M, F |
| Placenta | <LD | <LD | <LD | - | <LD | <LD | <LD | - |
| Cord Blood | <LD | <LD | <LD | - | <LD | <LD | <LD | - |
| Amniotic Fluid | <LD | <LD | <LD | - | <LD | <LD | <LD | - |
| Tissues | <LD | <LD | <LD | - | <LD | <LD | <LD | - |

dG, gestation day age; <LD below level of detection; Fetal tissues tested for ZIKV RNA were cortex (4 lobes), cerebellum, hippocampus, lung, liver, spleen, gonad, stomach, intestine, eye and thymus.

In the mid-gestation cohort, viremia was observed at 7 dpi for Dam 1M, and at 4 and 7 dpi for Dam 3M. Dam 2M did not have detectable viremia at any time point. ZIKV RNA was detected in saliva samples from Dam 2M on 7 dpi and Dam 3M at 4 and 7 dpi. Dam 1M did not have detectable ZIKV RNA in saliva at any time point. None of the dams had detectable ZIKV RNA in the vaginal swabs collected at any time point.

Reproductive tissues (vagina, cervix, uterus, ovaries) and lymph nodes (LNs) (axial, inguinal, mesenteric) were examined for ZIKV RNA from the dams at the time of necropsy. In the early-gestation cohort, of the four uterine sites sampled per dam, ZIKV RNA was detected in 1/4 in Dam 1E, 2/4 in 3E and in 3/4 samples in Dam 4E. Inguinal LN from Dam 1E was positive for ZIKV RNA and Inguinal and Axial LNs from Dam 4E were positive for ZIKV RNA. No ZIKV RNA was detected in ovaries, cervix or vaginal tissues for the early gestation dams.

None of the reproductive tissues collected from the mid-gestation cohort had detectable ZIKV RNA. Axial and Inguinal LNs from Dam 1M, Axial LNs from Dam 2M and 3M had detectable ZIKV RNA.

### Placental, amniotic fluid and fetal ZIKV Viral loads

Fetuses are coded to match the dams (eg. Dam 1 = Fetus 1). Placenta, fetal tissues and amniotic fluid collected at the time of necropsy from early and mid-gestation animals did not have detectable ZIKV RNA.

### ZIKV specific IgM and IgG

#### IgM

All dams were negative for ZIKV IgM prior to infection and remained negative through 7 dpi. Dams 1E, 2E and 3E from the early-gest cohort and Dams 1M and 2M from the mid-gestation cohort had IgM response from 14 through 42 dpi. Dam 3M had ZIKV IgM on 14 and 21 dpi and had the weakest IgM response compared to all other dams. Dams 1E and 3E from early-gestation and 1M from mid-gestation group exhibited the most robust IgM response (**Fig 2 A**).

**Figure 1.**
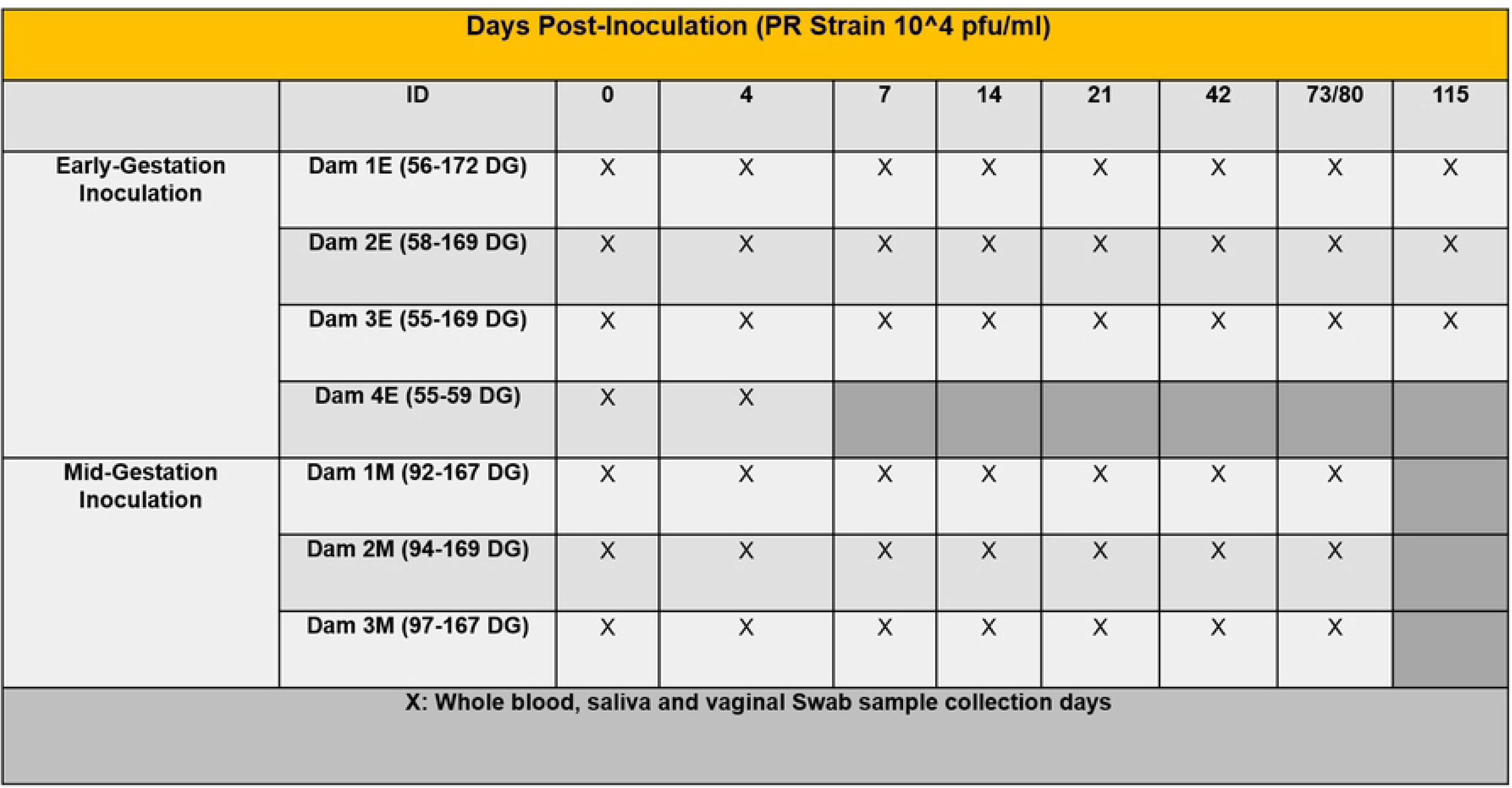
Schematic representation of the experimental design. Timed pregnant olive baboons at early gestation (n=4) and mid-gestation (n=3) were infected subcutaneously (1x10^4^ pfu, 1 ml volume, strain PRVABC59, 2015) on Day 0. Maternal blood, saliva and vaginal swabs were obtained on the indicated days for each animal. The gestational age at the time of infection ranged from 55-172 days gestation (dG) for early and 92-169 dG for mid-gestation animals. Dams were euthanized at the end of study for collection of maternal and fetal tissues except for Dam 4 euthanized at 59 dG after fetus was aborted 5 days post infection (dpi). 3 timed pregnant control dams were euthanized at 166, 169 and 171 dG.

**Figure 2.**
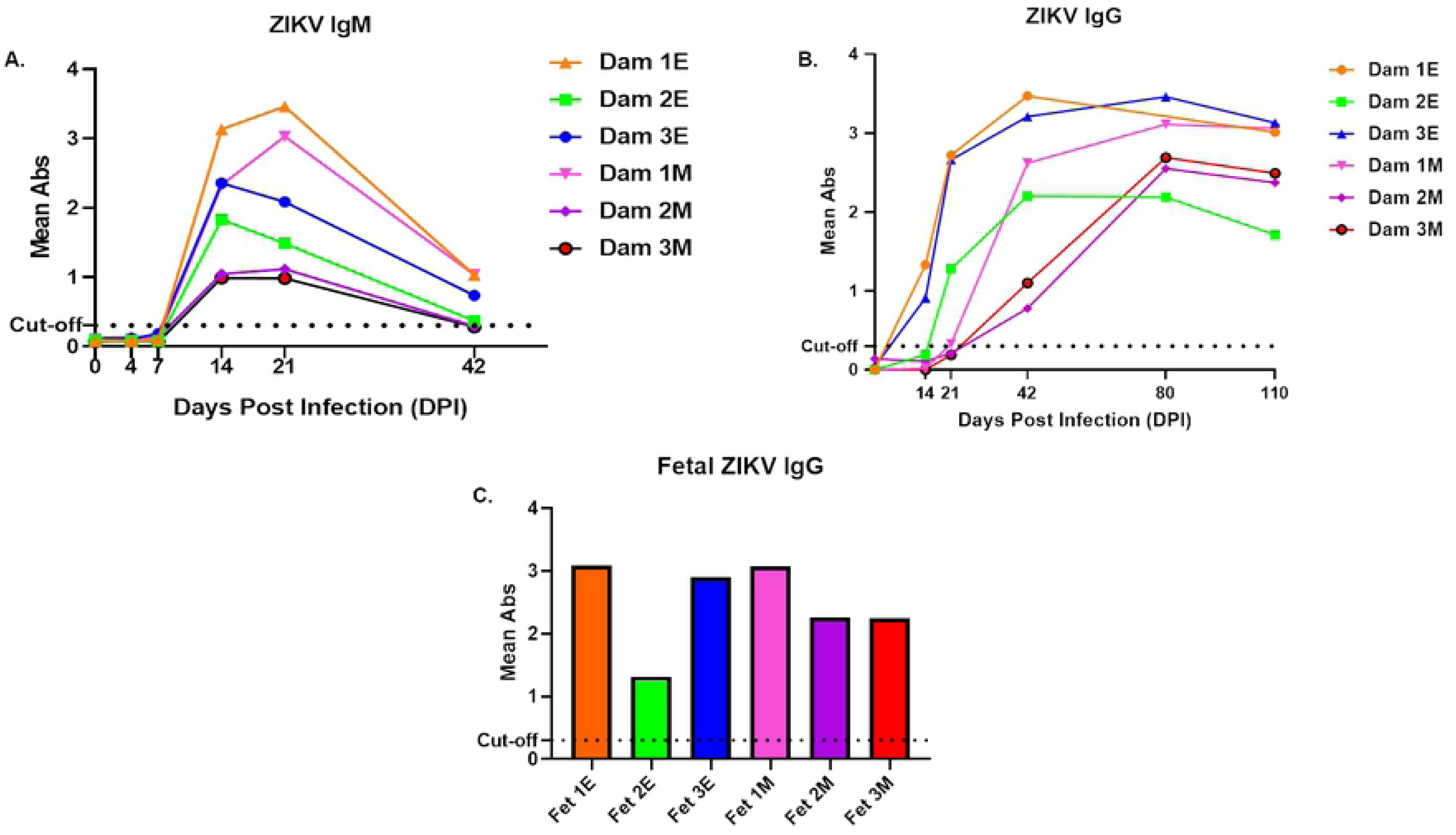
Detection of anti-ZIKV antibody responses in pregnant baboon serum. Antibodies against ZIKV was determined by ELISA for maternal IgM (**A**) and IgG (**B**) and fetal IgG (**C**). anti-ZIKV IgM were detected at 14 days post-infection in all six baboons sampled at this time point. In the early-gestation group, dams 1E and 3E had ZIKV IgG at 14 dpi and dam 2E had at 21 dpi. In the mid-gestation group, all three dams had IgG by 21dpi. All six fetuses had anti-ZIKV IgG at levels similar to the mothers indicating efficient FcγR transfer of IgG across the placenta.

#### IgG

No dams had serum ZIKV IgG by 7 dpi. In the early-gestation cohort, Dams 1E and 3E had ZIKV IgG from14 dpi, gradually increasing to 115 dpi. Dam 2E had ZIKV IgG from 21-110 dpi. In the mid-gestation cohort, Dam 1M had increasing ZIKV IgG from 14 to 73 dpi and Dams 2M and 3M had ZIKV IgG from 21 to 73 dpi. ZIKV IgG was also detected in the cord blood of all fetuses from early and mid-gestation dams (**Fig 2** **B, C**).

### CNS Histology and Immunohistochemistry

#### Brain pathology

Post necropsy, fetal brains were assessed for gross structural defects, primarily, overall size and gyri and sulci formation. Fetal brains from ZIKV infected dams exhibited a range of irregularities of the major gyri and sulci in the different lobes of the cerebral cortex regardless of gestation age at infection. Fetuses 1E, 3E (early-gestation cohort), 1M and 3M (mid-gestation cohort) had the most visibly malformed gyration and sulcation (**Fig 3**). The brain from Fetus 2M (mid-gestation) was similar in appearance to control brains. Malformations highlighted in **Fig 3** include visible absence of prominent gyris and sulcis, enlarged gyris and abnormal sulcal length and depth. Pathology was also observed in the cerebellum of fetus 3E (early-gestation) and 3M (mid-gestation) with reduced cerebellar size compared to the control brains (**Fig 3**).

**Figure 3.**
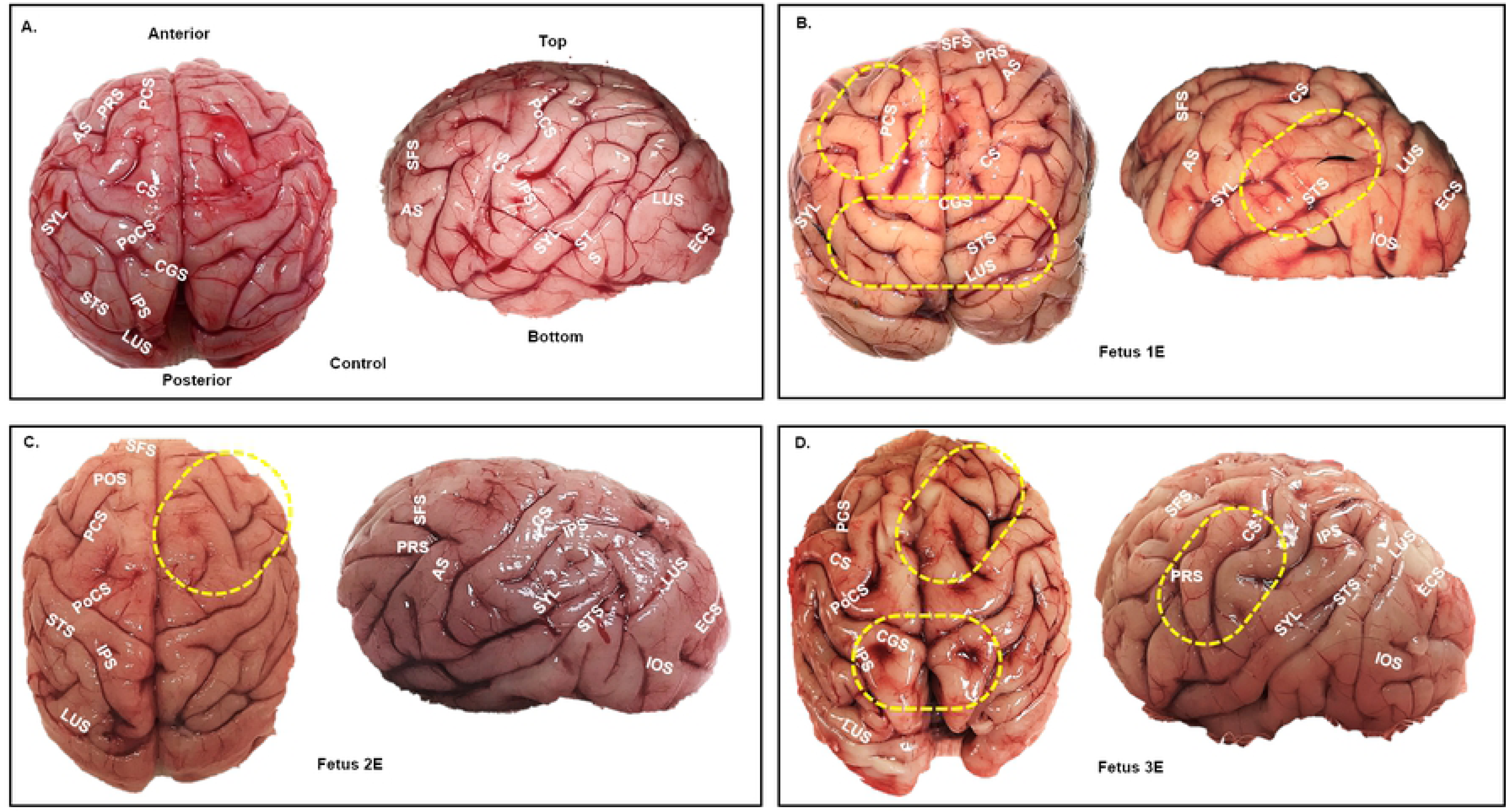
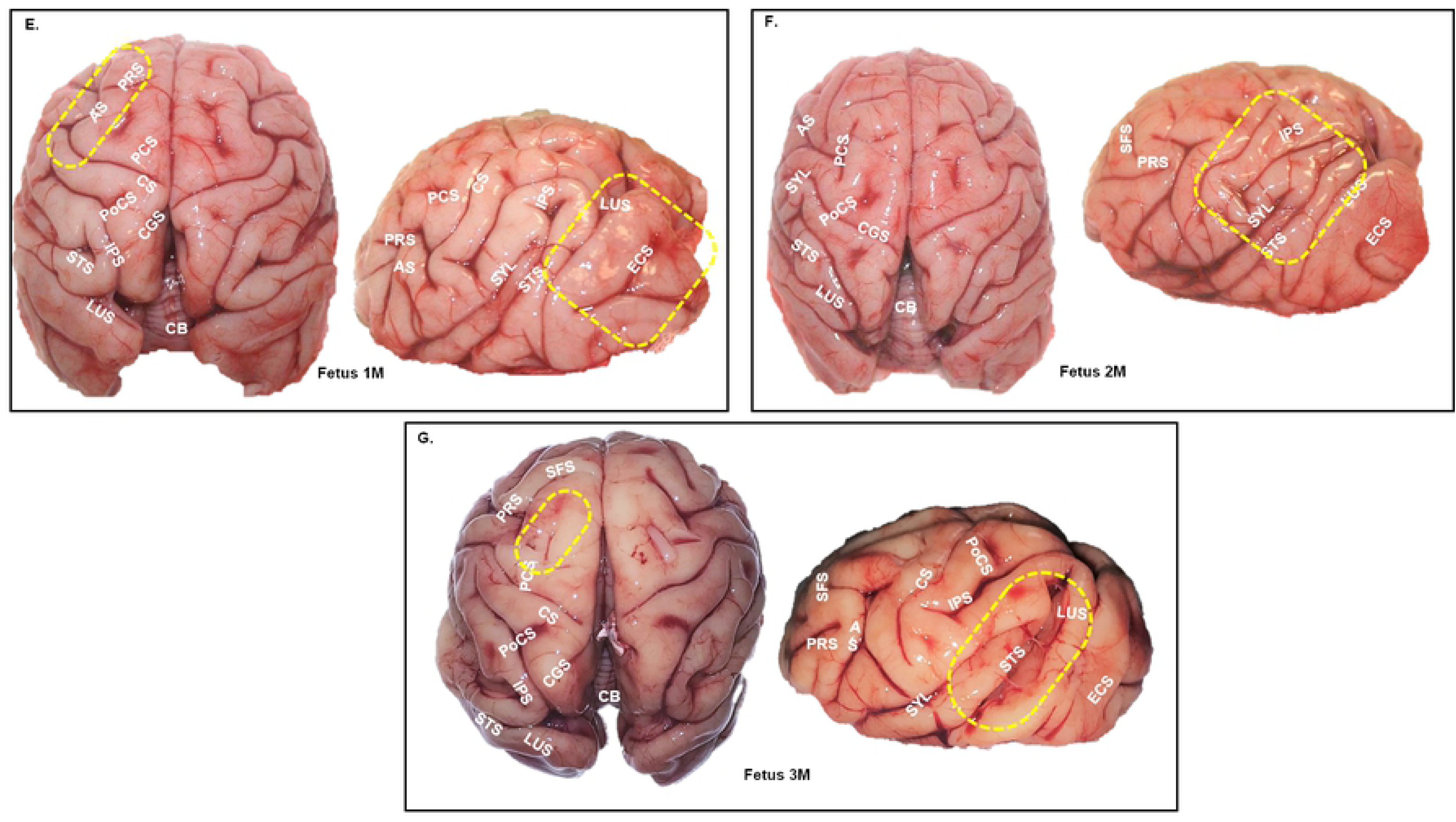
Gyri/sulci malformations in the early and mid-gestation ZIKV brains compared to the control brain (**A**). Top and side view of fetal brains showing major gyri and sulci formations. Areas marked by yellow dotted ovals highlight significant gyral/sulcal malformations in various areas of the cortex. Malformations were mainly characterized by absence of gyri/sulci, enlarged gyri, abnormal sulcal length and depth and asymmetric gyri/sulci malformation across the cerebral hemispheres compared to the control fetal brain **(A)**. Early-gestation infected fetuses **(B, C, D)**; Mid-gestation infected fetuses (**E, F, G**). Arcuate Sulcus (AS), Cerebellum (CB), Central Sulcus (CS), Cingulate Sulcus (CGS), Entocalcarine Sulcus (ECS), Inferior Occipital Sulcus (IOS), Inferior Parietal Sulcus (IPS), Lunate Sulcus (LUS), Parieto Occipital Sulcus (POS), Pre-Central Sulcus (PCS), Post Central Sulcus (PoCS), Principal Sulcus (PRS), Superior Frontal Sulcus (SFS), Superior Temporal Sulcus (STS), Sylvian Fissure/Lateral Sulcus (SYL).

#### Histology of Frontal Cortex and Cerebellum

H&E staining of the frontal cortices of ZIKV fetuses showed a general disorganization of the six-layered cortex in early (1E, 2E, 3E) and mid (1M, 2M, 3M) gestation compared to the control fetal brains. The category and degree of disorganization of the fetal cortex in different layers of the cortex varied among the fetuses (**Fig 4 A**). Histology of F1E, F2E and F3E (early gestation infection) and F1M (mid gestation infection) cortical plate (CP) showed disorganization of the neuronal population in layer IV and V with F3E CP showing substantial lack of layer V specific pyramidal cells. Fetuses 1E, 2E, 3E (early) and 1M (mid) exhibited focal hemorrhage within the CP. The CP of mid-gestation fetus F2M showed disruption in layer specific cell populations including the presence of pyramidal cells throughout all layers. Layer V of the CP of mid-gestation fetus F3M exhibited considerably fewer pyramidal cells and focal hemorrhage throughout the CP (**Fig 4 B**).

**Figure 4.**
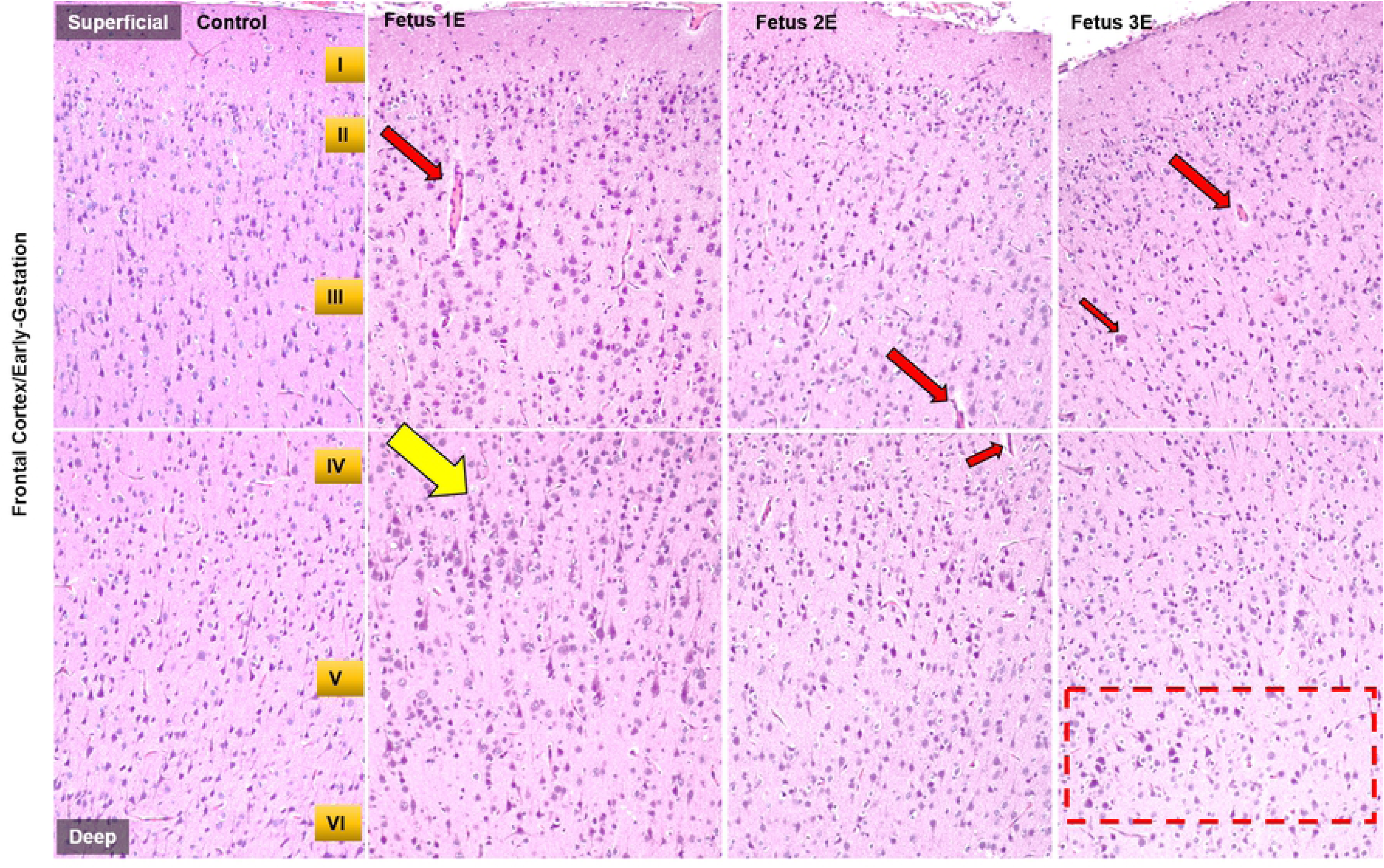
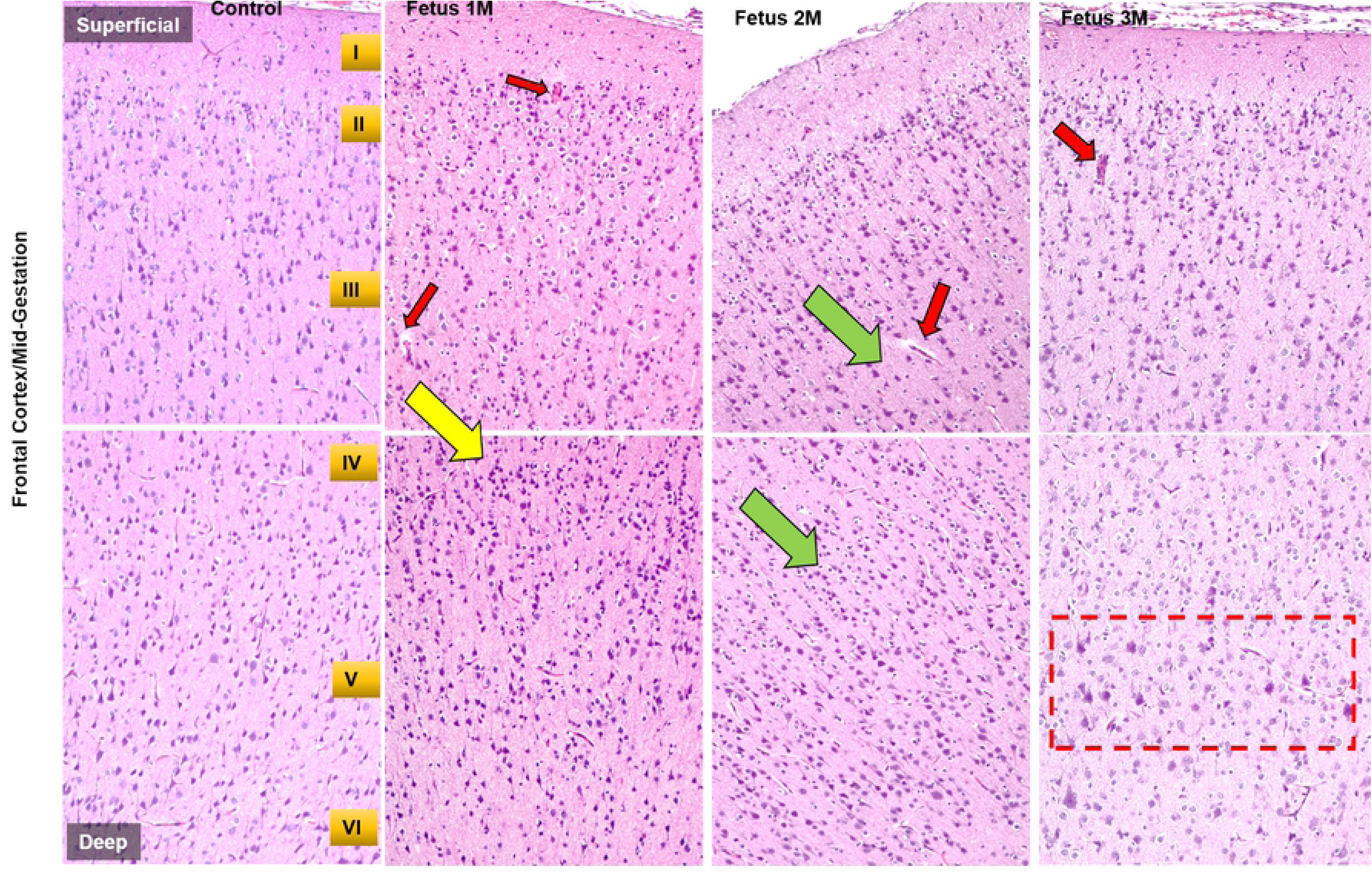
ZIKV infection disrupts formation of six-layered cortex in a developing fetal brain. Representative images from H&E staining of control and early-gestation (**A**) and mid-gestation (**B**) cortical plate of the fetal frontal cortex. The histological stain shows control fetal brain with orderly formation of six layers of cortex compared to the disturbance of this process in the (**A**) early and (**B**) mid-gestation fetal cortical plates. Yellow arrows represent disorganization in different layers, green arrows represent undefined layers and its cell population, red arrows represent focal hemorrhage and vasculitis and red dotted rectangle represent substantial lack of pyramidal cells in Layer V of the cortical plate. (bar = 100μm)

H&E staining of the cerebellum showed a marked disruption of structural integrity of cerebellum of ZIKV fetuses compared to the control (**Fig 5**). In the early-gestation cohort, Fetuses 2E and 3E showed the most cerebellar defects including loss and/or distortion of Purkinje cells in the ganglionic layer of the cerebellum and focal hemorrhage in the granular layer compared to the control cerebellum. The cerebellum of F1E had the fewest observable deficits. The cerebellum of F3E had the greatest Purkinje cell layer damage coupled with hemorrhage.

**Figure 5.**
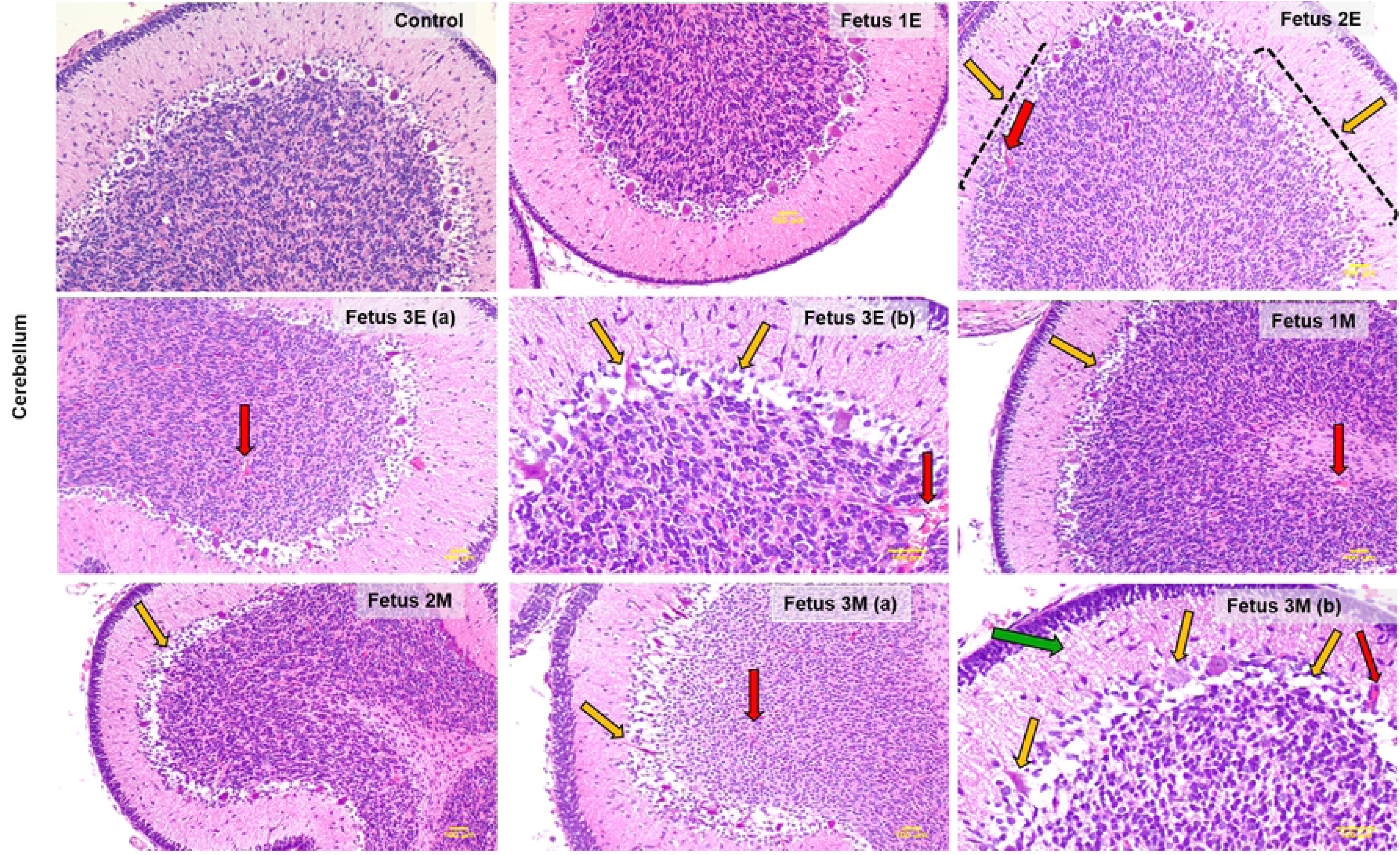
ZIKV infection causes structural cerebellar damage in early and mid-gestation infected fetal brain compared to the control. Representative images from H&E staining of control, early (**Fetus 1E, 2E, 3E**) and mid-gestation (**Fetus 1M, 2M, 3M**) cerebellum showing damaged and absent purkinje cells in the ganglionic layer as marked by the yellow arrows. The red arrows mark focal hemorrhage and vasculitis in the different layers of the cerebellum and the green arrow marks the loss of cell volume in the molecular layer of fetus 3M (**7b**). All images taken at 4x magnification except 3Eb and 3Mb which are 10x magnification to highlight severity of the cerebellar damage in Fetus 3E and 3M. (bar = 100μm)

In the mid-gestation cohort, the cerebellum of F1M, 2M and 3M exhibited damage to Purkinje cells and focal hemorrhage in different layers of the cerebellum. Fetus 3M showed the most damage to the cerebellar structure with marked loss of Purkinje cells in the ganglionic layer, loss of cell populations in the molecular layer and extensive hemorrhage in the granular layer of the cerebellum.

### Gliosis

#### Microglia

In the frontal cortex, microglia (Iba-1) were counted in the white matter of each section. Early-gest infected fetal frontal cortices exhibited a significant increase in Iba-1+ cells in the white matter compared to the control brains (p=0.03). Mid-gest fetal frontal cortices did not exhibit increased Iba-1 immunoreactive microglial population compared to the control fetuses (**Fig 6 A**). In the hippocampus, Iba-1+ microglial cells were counted in the dentate gyrus (DG). No difference in the number of Iba-1+ cells in the DG between ZIKV infected and control fetuses was seen (data not shown).

**Figure 6.**
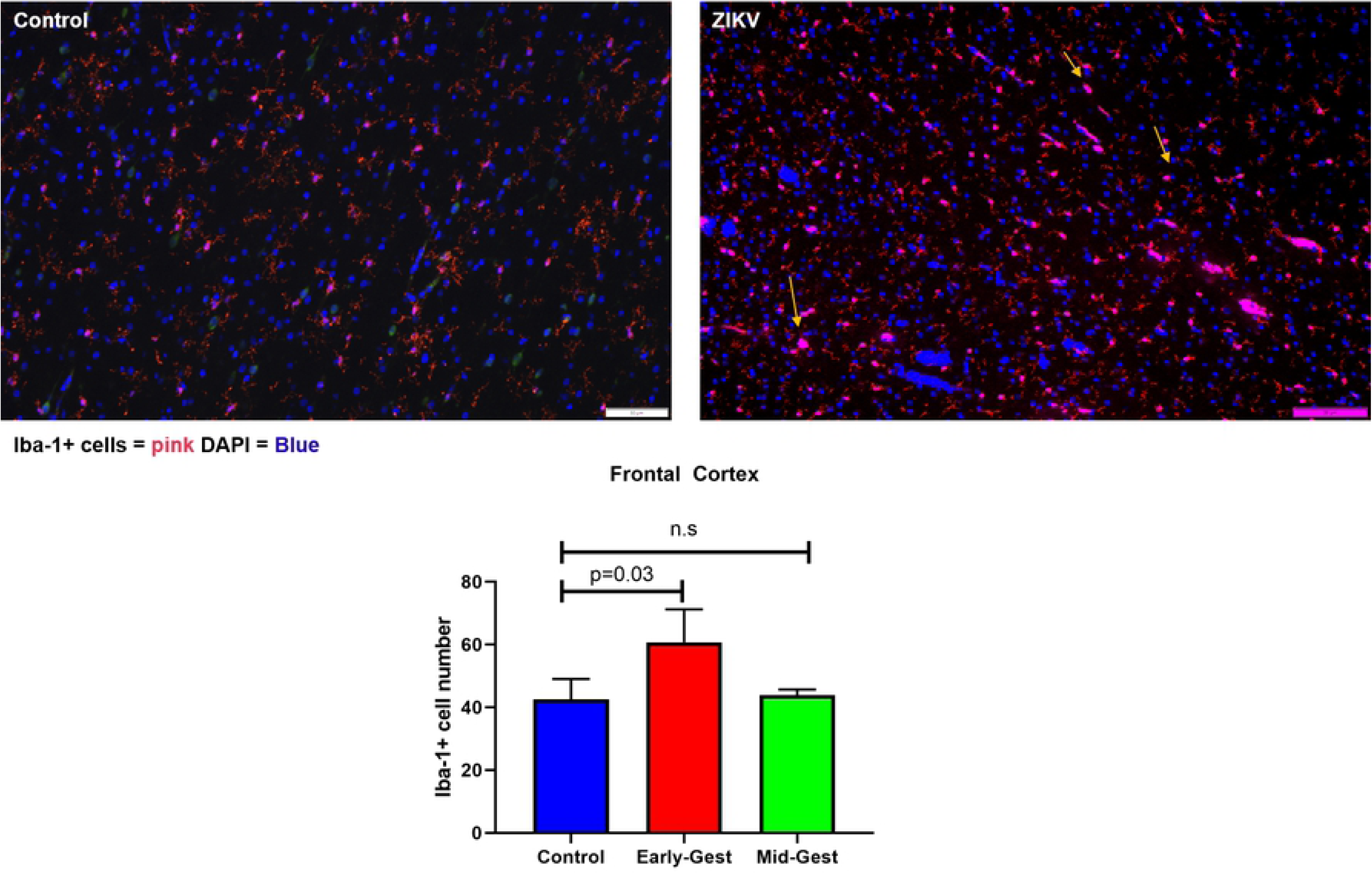
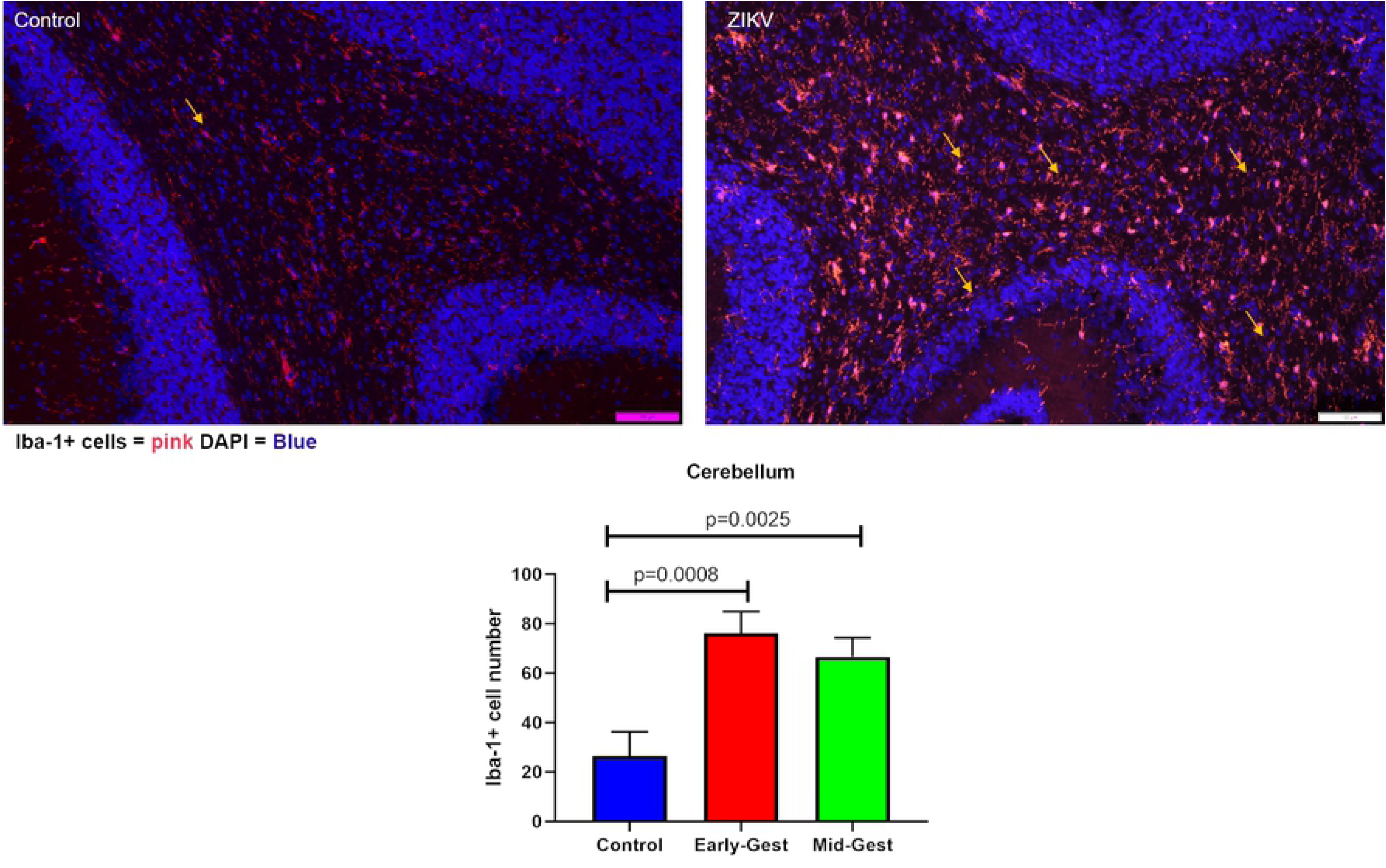
Immunofluorescence (IF) for neuroinflammatory marker for microglia (Iba-1) in the frontal cortex (**A**), and cerebellum (**B**) of control and ZIKV fetuses. In the frontal cortex (**A**) white matter, significant increase in Iba-1+ cells in number and staining intensity signaling reactive microglial population were observed as marked by yellow arrows in the early but not in mid-gestation infected brains compared to the control. In the cerebellum (**B**), both early and mid-gestation infected brains had significantly more Iba-1+ reactive microglia in the granular layer compared to the control. (bar = 50μm; mean ± SEM for four sections per fetus).

In the cerebellum, a significant increase in immunoreactive microglial cells was observed suggesting marked cerebellar inflammation in both the early and mid-gestation fetuses compared to the control (**Fig 6 B**). The early-gestation ZIKV infected fetal cerebellum exhibited 4-fold increase in Iba-1+ microglial cells (p=0.0008) and the mid-gestation ZIKV infected fetuses exhibited 3-fold increase in Iba-1 immunoreactive microglial cell population in the cerebellum compared to the control cerebellum (p=0.0025).

#### Astrocytes

In the frontal cortex, there was a significant increase in GFAP+ cells in both early and mid-gestation infected fetal brains compared to control (**Fig 7**). Significant increases in the astroglial cell population in the fetal brain at term is a mark of astrogliosis or reactive gliosis. Among the gestation groups, the early-gestation infection fetal cortex had significantly more GFAP positive cells compared to the control (p=0.0035) than the mid-gestation group (p=0.058). We also observed not only astrocytic hyperplasia but also hypertrophy, increased GFAP marker intensity and higher number of cytoplasmic processes suggesting changes in the astrocytic morphology and expression of cytoskeleton proteins which are the hallmarks of reactive gliosis. There were no apparent changes in the hippocampal or cerebellar astrocytic populations in either early or mid-gestation infected fetuses despite the noted histological changes observed above.

**Figure 7.**
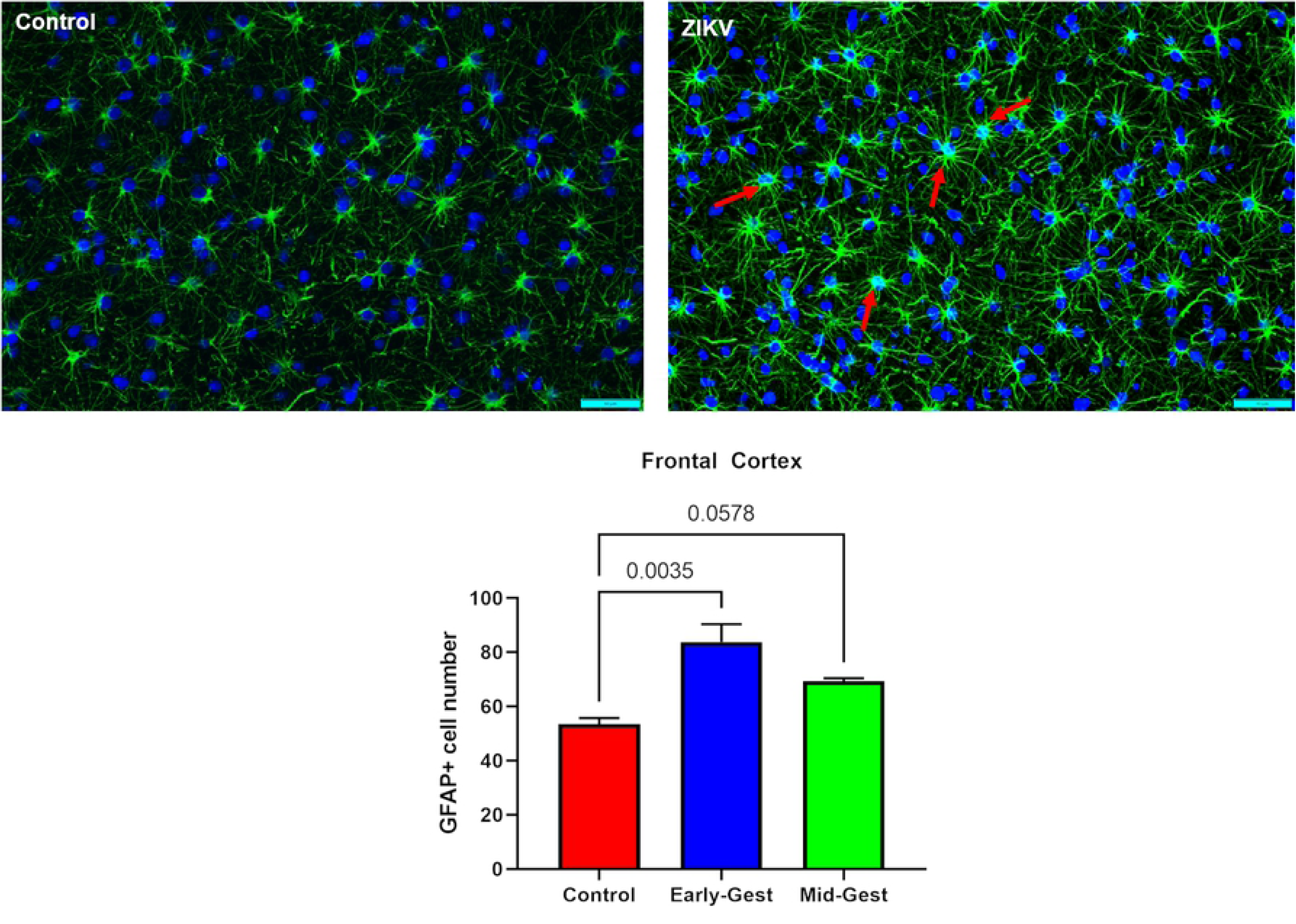
Immunofluorescence (IF) for GFAP representing astrogliosis in the frontal cortex of early and mid-gestation ZIKV infected fetal baboons compared to the control brain. In both early and mid-gestation infected frontal cortex, significant increase in GFAP+ cells with increased GFAP staining intensity and morphological changes such as longer and more ramified cellular processes and larger cell bodies were seen compared to the control brain. (bar = 50μm; mean ± SEM)

### Olig-2 (Oligodendrocyte precursor)

We evaluated the effects of ZIKV infection on the oligodendrocyte cell population in the cerebellum using Olig-2 immunostaining as a marker for oligodendrocyte precursor cell population which matures to become oligodendrocytes. Olig-2 immunostaining showed a significant decrease in the oligodendrocyte precursor cell population in the cerebellar white matter of ZIKV infected fetuses compared to the control cerebellum (**Fig 8**). Early-gest cerebellum showed significant reduction in Olig-2 immunostained cells compared to the control (p=0.005) whereas the mid-gest fetal cerebellar showed less but not significant reduction in Olig-2+ population (p=0.16) compared to the control fetuses. Compared to the cerebellum, no significant difference in Olig-2+ cell population was seen in the frontal cortices of ZIKV fetuses of both gestation ages compared to the control (data not shown).

**Figure 8.**
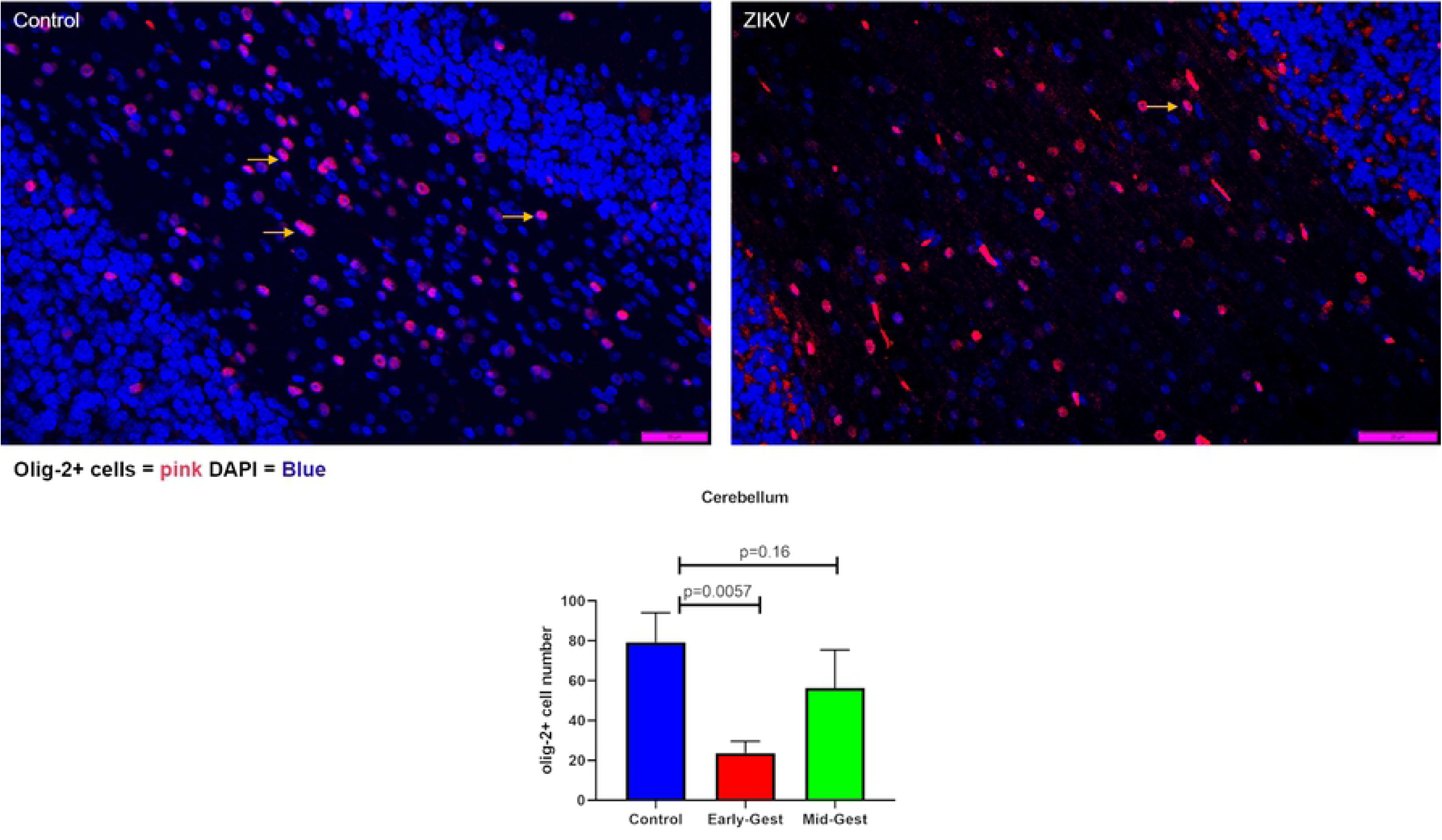
Immunofluorescence (IF) for Olig-2 (oligodendrocyte precursor) in the granular layer of the cerebellum. In the control cerebellum, numerous Olig-2+ cells marked by yellow arrows could be seen in the granular layer whereas in the early-gestation infected ZIKV cerebellum, significant decrease in Olig-2+ cell population was noted. However, mid-gestation infected fetal cerebellum did not show difference in Olig-2+ cell population compared to the control. (bar = 50μm; mean ± SEM)

### Nestin and Doublecortin (DCX) in the Hippocampus

In order to determine the neurological impact of ZIKV infection on neural progenitor cells (NPCs) and newly differentiated and migrating immature neurons, we immunostained control and ZIKV infected hippocampal sections using Nestin and Ki-67 (NPCs; mitotic cells) and Doublecortin (DCX, differentiation and migration) as markers. Hippocampal sub granular zone (SGZ) in the dentate gyrus (DG) is a neurogenic niche in all stages of brain development pre- and post-birth and throughout life. Hence, active neurogenesis takes place in the DG of the hippocampus with generation of NPCs as well as differentiation, migration and survival of the newly formed neurons.

Neurite length of Nestin positive cells in the DG were reduced in the ZIKV infected fetuses compared to the control hippocampus (p=0.09, **Fig 9 B**). The general appearance of neurites of the Nestin immunoreactive NPCs in the DG looked disorganized with loss or distortion of neurites and uneven distribution of Nestin^+^ cells along the DG compared to the orderly distribution of symmetrical and long tracks of neurites of Nestin immunoreactive cells in control DG (**Fig 9 A**). No difference in actively replicating progenitor cell population (Ki-67/Nestin+ cells) was seen between the ZIKV infected and control hippocampus (**Fig 9 C**).

**Figure 9.**
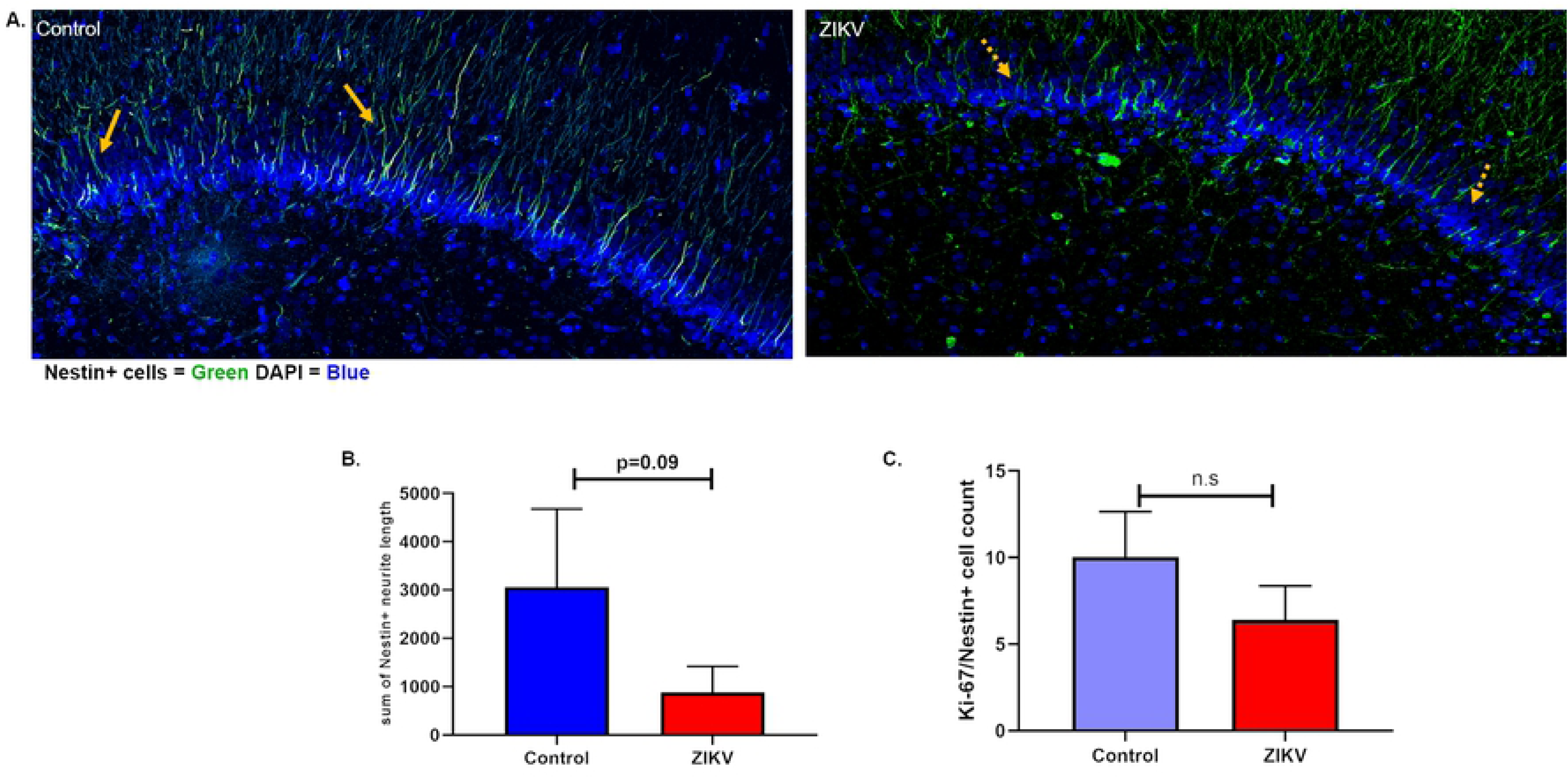
Immunofluorescence (IF) for neural progenitor cell (NPC) marker Nestin in the sub granular zone (SGZ) of the hippocampal dentate gyrus (DG). (**B**) Neurite length in the Nestin+ cells were reduced in the DG of ZIKV infected fetal hippocampus with the appearance of degeneration as compared to the control hippocampus where neuronal processes appeared uniform and abundant as marked by orange arrows. (**A**) Besides shorter and degenerated neurite processes in Nestin+ cells in the DG, disorganized distribution of these cells along the DG compared to the control can also be seen. (**C**) No difference in actively dividing progenitor cells were seen in the DG between ZIKV and control hippocampi. (bar = 50μm; mean ± SEM)

DCX immunostaining of ZIKV and control hippocampi showed decrease in DCX+ staining intensity in the DG of ZIKV infected fetuses compared to the control but not reaching statistical significance (**Fig 10 B**). DCX+ cells in the DG in the control hippocampi showed robust expression of the marker in the nucleus, soma and the neurite processes whereas in the ZIKV infected fetal hippocampus, there were less DCX immunostaining, and shorter and disorganized neuronal processes (**Fig 10 A**).

**Figure 10.**
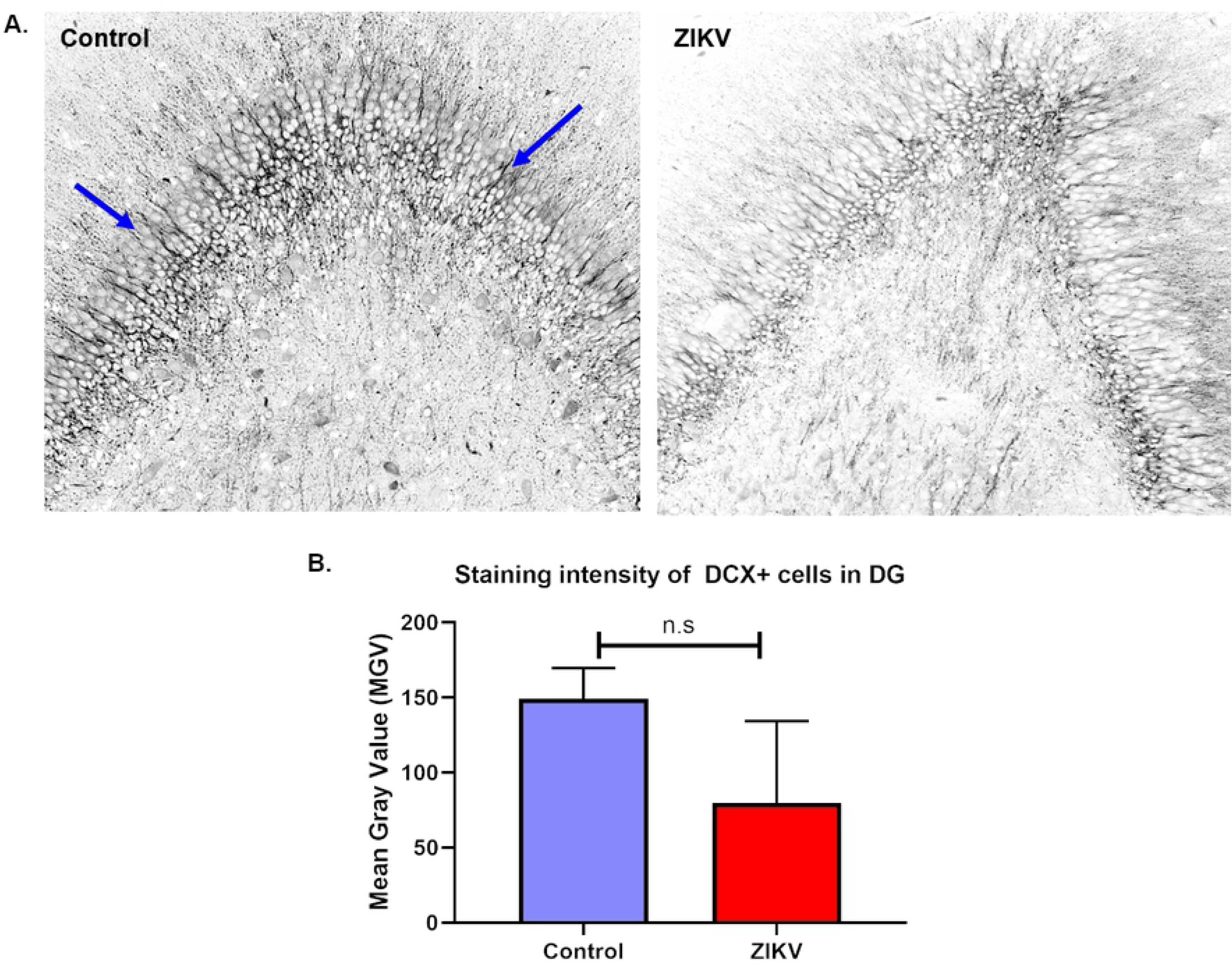
Immunofluorescence (IF) for neuronal migration marker doublecortin (DCX) in the dentate gyrus (DG) of ZIKV infected and control hippocampus (**A**). Staining intensity of DCX+ neurons localized in the DG was measured as mean grey value (MGV). The control DG showed uniform distribution of DCX+ neurons with long, uniform and abundant neuronal processes as marked by arrows whereas the ZIKV DG had disorganized population of DCX+ cells with lack of neuronal processes and some with degeneration. (**B**) Difference in MGV of DCX+ neurons between ZIKV and control hippocampus was not statistically significant. (bar = 50μm, mean ± SEM)

### Placental and Uterine Histology and IF

The histopathology of ZIKV infection in our early and mid-gestation infection groups showed pathological changes compared to the control (**Fig 11**). In control placenta, the chorionic villi were seen as mature late gestation villi with an intact syncytiotrophoblast (STB) layer, multiple unobstructed fetal capillaries and few fibrin and calcium deposits throughout the villi on the fetal side (**Fig 11 A**, **B**). In ZIKV groups, placental pathology notably observed were disruption of STB layers and unusual STB formation (**Fig 11** **E, G),** delayed villous maturation characterized by presence of more immature villi with reduced branching and less terminal villi intermixed with intermediate villi compared to the control placenta (**Fig 11 D**). ZIKV placentas also had more avascular villi (villi without fetal capillaries) (**Fig 11 D**), partially or fully thrombosed vessels with calcium mineralization and fibrin deposits (**Fig 11** **E, F, G**) and thrombosed fetal blood vessels in the chorionic plate undergoing mineralization and surrounded by fibrin deposits (**Fig 11 H**). Dam 1E had significant hemorrhage in the intervillous space (**Fig 11 C**). Overall, early-gestation infected dams had more placental pathology resulting from ZIKV infection compared to mid-gestation infected dams.

**Figure 11.**
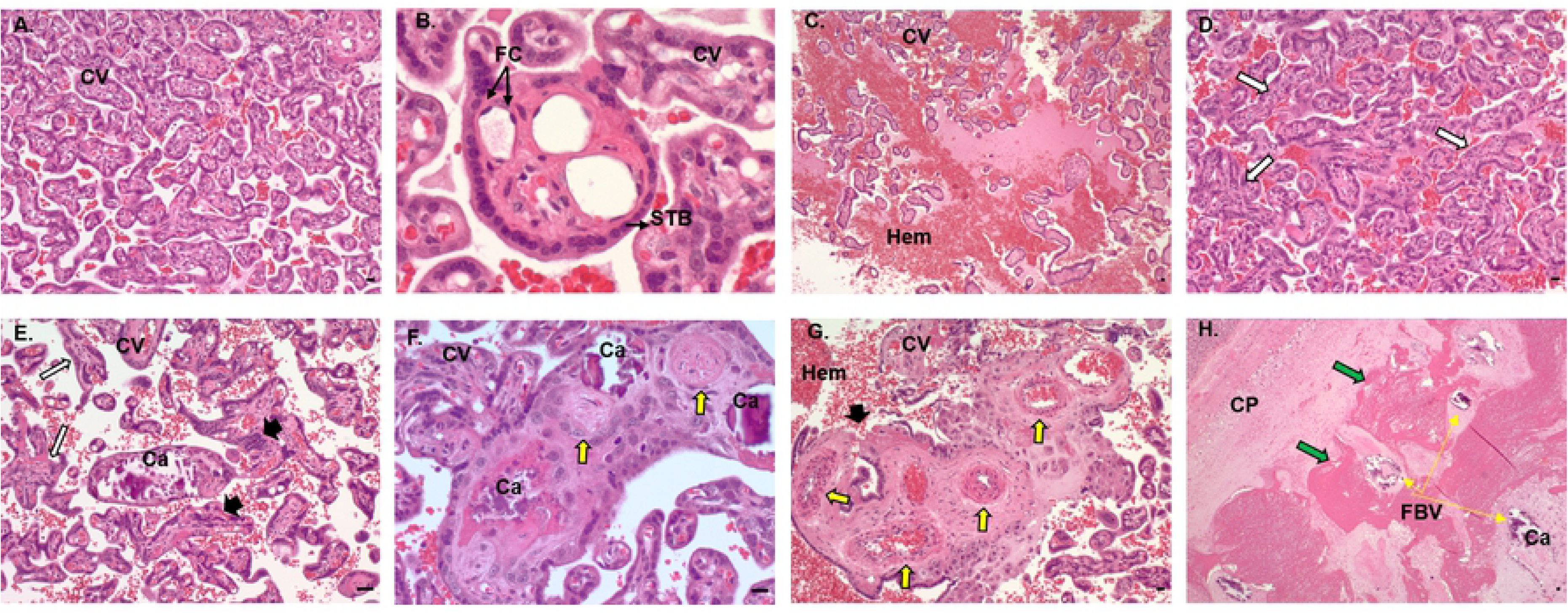
Histopathology of control and ZIKV infected placenta. (**A, B**) Control placenta with normal chorionic villi structure with uninterrupted syncytiotrophoblast layer and numerous fetal capillaries lined with fetal endothelial cells. (**C, H**) Placenta from ZIKV infected dams showing multiple changes in the villi and vascular structure such as immature and avascular villi (white arrows), disrupted and abnormal syncytiotrophoblast layer (black arrowheads), fetal capillary thickening, thrombosis and mineralization (yellow arrows) and fibrin deposits (green arrows) surrounding fetal vessels undergoing thrombosis and mineralization. CV, chorionic villi; STB, syncytiotrophoblasts; FC, fetal capillaries; Hem, hemorrhage; Ca, calcification; CP, chorionic plate; FBV, fetal blood vessels. (bar = 50μm).

Placental sections from early and mid-gestation dams exhibited increased MAC387 (macrophage) positive cell populations in the villous, intervillous space (IVS) on the fetal and maternal side compared to control (**Fig 12 A**). One distinction between MAC387+ cells in the early vs mid-gestation placenta was the location of these cells in the two groups. In the early-gestation placenta (Dam 1E, 2E, 3E), the MAC387+ cells were found interspersed in the terminal villi and IVS whereas in the mid-gestation placenta (Dam 1M, 2M and 3M) the MAC387+ cells were concentrated in the chorionic plate than villi on the fetal side to a greater degree (**Fig 12 A**).

**Figure 12.**
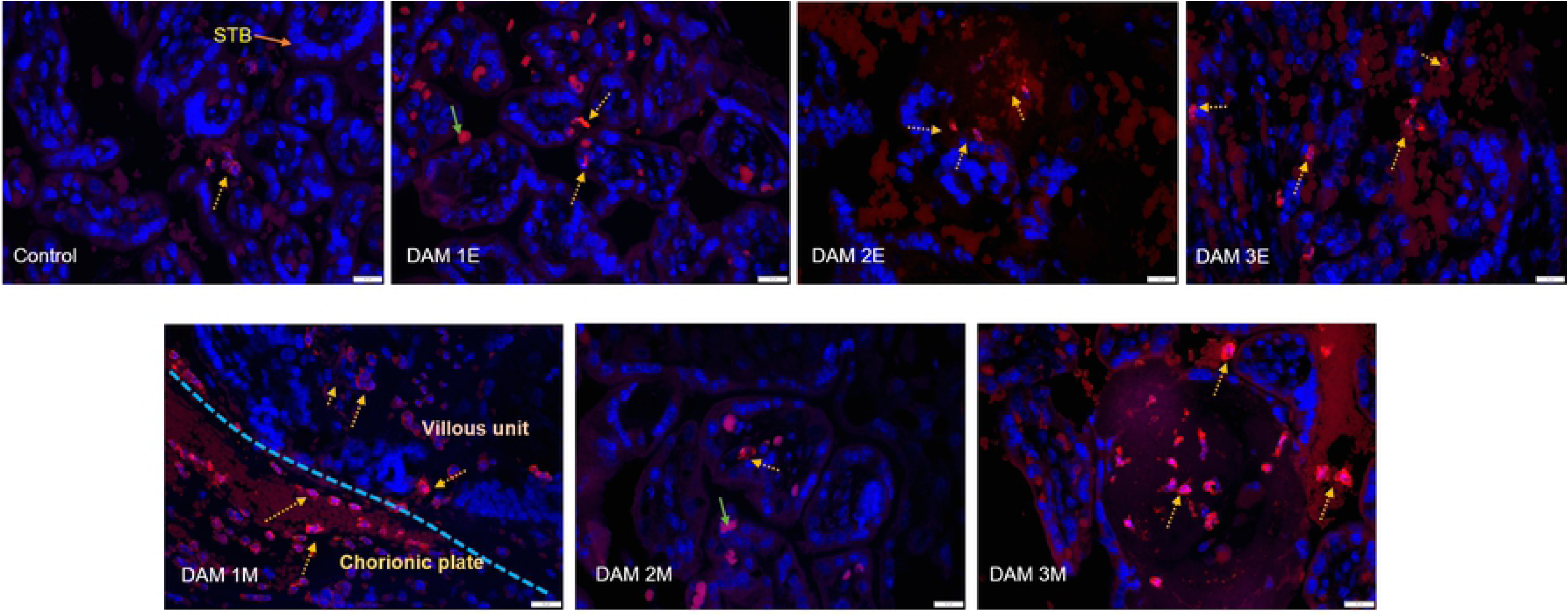
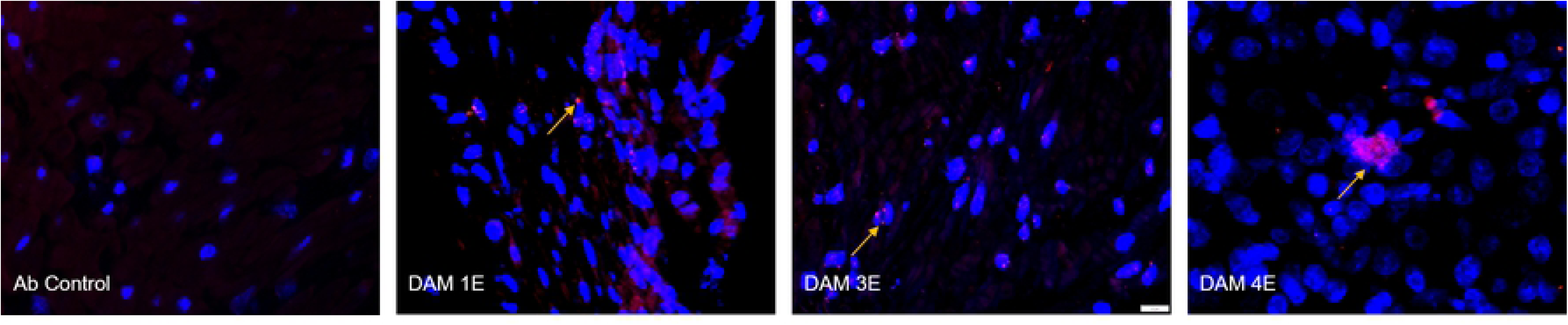
Immunofluorescence (IF) for placental macrophages (MAC387) and ZIKV (pan flavivirus) in the uterus. (**A**) Macrophage (red; DAPI: blue) staining in the placenta indicated with dotted yellow arrows was seen mostly in the villous and intervillous space in the placenta in Dams 1E, 2E, 3E, 2M and 3M and in chorionic plate on the fetal side in dam 1M (blue dotted line separating the fetal and villous sides). Only occasional macrophages were observed in Dam 1E placenta and abundant macrophages were observed in Dam 3M within villi compared to the control placenta (green arrows denote auto-fluorescing red blood cells). (bar = 20µm) (**B**) Pan-flavivirus immunofluorescence (Red:ZIKV; blue:DAPI) staining in the uterus. ZIKV IF was observed in the endometrial stromal cells in Dams 1E, 3E and 4E (yellow arrows). Dam 4E had widespread ZIKV IF staining with clusters of ZIKV infected cells. None of the mid-gestation dams showed ZIKV IF staining in the uterus. (bar = 20µm)

Three out of four early-gestation dams (Dams 1E, 3E and 4E) were positive for ZIKV IF in the uterus (**Fig 12 B**). Uterus of these dams also tested positive for ZIKV RNA by q-PCR (**Table 1**). ZIKV IF was detected primarily in the endometrial stromal cells. In the case of Dam 4E, ZIKV IF was seen widespread and more diffused in the endometrial stromal cells. Dam 4E also aborted 5 days post infection. Dams in the mid-gestation cohort were negative for ZIKV IF in the uterus.

## DISCUSSION

The 2015 ZIKV epidemic in Brazil revealed devastating clinical consequences of ZIKV infection to the fetus during pregnancy as what we now refer to as congenital zika syndrome (CZS). CZS includes a wide array of neurological and developmental disorders in the fetus and the infant post-birth. Therefore, studying ZIKV and its pathophysiology remains crucial in understanding how viruses like ZIKV cross the placental barrier to infect the developing fetus, causing permanent damage to the developing brain. Studies using mouse and non-human primate models such as rhesus macaques have shown some neuropathology, primarily region specific microcalcifications, hemorrhage and gliosis but no overt microcephaly, inflammation, cortical malformations and structural changes such as cerebral gyri/sulci formations and cerebellar atrophy. We described previously how subcutaneous ZIKV infection in the olive baboons at mid-gestation with a relatively modest dose (1x10^4^ ffu) of the French Polynesian isolate resulted in vertical transfer of the virus to the fetus associated with fetal demise in one pregnancy (14 dpi) and significant fetal CNS pathology in a second pregnancy (21 dpi) [28]. To our knowledge, this is the earliest cortical neuropathology post-ZIKV infection in a primate in a viable fetus that was not aborted or underwent *in utero* death, and the first description of loss of radial glia (RG) and RG fibers in primates in response to ZIKV. Besides the loss of RG fibers, we also observed astrogliosis, loss of oligodendrocyte precursor cells and indices of neuroinflammation with increased Iba-1 (microglia) and IL-6 (proinflammatory cytokine) immunostaining in the 21-dpi fetal cortex. Since the acute study helped define the timeline of fetal vertical transmission and neurological damage after ZIKV infection during mid-gestation in our baboon model, in the present study we examined the longer-term pathological outcome in term gestation fetuses as a result from ZIKV infection during early and mid-gestation. Early and mid-gestational ages are crucial to study the impact of ZIKV infection during pregnancy since a more severe CZS phenotype is observed clinically in human fetuses exposed to ZIKV in mothers infected earlier vs later in gestation [4]. Our findings show severe ZIKV induced CZS in fetuses from dams infected early and mid-gestation consistent with CZS reported in human fetuses due to ZIKV infection.

Following ZIKV infection, all pregnant dams in early-gestation (n=4) and 2/3 mid-gestation (n=3) exhibited viremia within the first week post-infection (4-7 dpi). Viremia at any other time points up to the day of necropsy was not observed in any of the pregnant dams unlike the pregnant macaques where prolonged (or re-emergent) viremia for several weeks up to 70 dpi has been routinely reported [31, 32, 35–37]. While prolonged viremia (46 to 53 days) has been reported in pregnant women [51], it should be noted that this was restricted to five cases after a search of the entire U.S. Zika Pregnancy Registry and as such, prolonged viremia during pregnancy in women appears rare and no correlation between prolonged viremia and increased incidence of CZS was found. vRNA was also detected in saliva samples from pregnant dams in early (3/4) and mid-gestation (2/3) groups. vRNA was not detected in vaginal swab samples from early and mid-gestation dams except Dam 3 (early-gest) where samples from days 4, 7, 14 and 21 post infection tested positive for ZIKV RNA. All the dams had timely and robust IgM and IgG response to ZIKV infection with efficient transfer of ZIKV IgG to the fetal compartment at term gestation.

vRNA was not detected in near term placental samples, cord blood, amniotic fluid and fetal tissues collected during necropsy from early and mid-gestation groups. However, vRNA persisted in the various lymph nodes of one early gestation and three mid-gestation infection dams. While two of the early-gest infected had persistent and high ZIKV viral load in uterine tissue samples collected at term gestation, none of mid-gestation dams exhibited uterine ZIKV RNA. As such, the early gestation uterus is a potential harbor for ZIKV, and the observation that none of the mid-gestation dams exhibited ZIKV RNA in uterus suggests a change in viral susceptibility as gestation progresses in the uterus.

Detection of ZIKV in near term uterus and lymph nodes in pregnant dams infected early-gestation but not mid-gestation may explain why early-gestation infections have a more severe fetal outcome compared to later gestational ages of infection. It is possible that the uterus and lymph nodes were serving as viral reservoirs throughout gestation. We cannot answer if viral presence in the uterus was persistent throughout the pregnancy in these dams or the virus re-emerged from lymph nodes serving as viral reservoirs in the uterus later in gestation. It is unknown if infectious virus was present at term in uterus or lymph nodes in these Dams but considering the lability of RNA, it is likely that live virus did persist for some time, and as IF demonstrated in the stromal layer, providing contact with the fetal membranes. We do not know the mechanism by which ZIKV evaded the maternal immune system and what role the continued presence of ZIKV in reproductive tissues and lymph nodes play in fetal development and future infections. IF staining for a macrophage marker MAC387 showed that ZIKV infection regardless of gestation age increased MAC387+ cell population in both the maternal and fetal sides of the placenta even in the tissues from dams with undetectable ZIKV RNA compared to the control tissues suggesting placental inflammation due to ZIKV infection.

The placenta plays a critical central role in fetal development and outcome. One of the questions still unanswered is the role placental pathology such as vascular insufficiency and inflammation due to ZIKV infection in vertical transmission and fetal outcome. Studies in pregnant macaques have shown a range of placental pathology resulting from ZIKV infection ranging from no discernable pathology to mild through severe pathology including deciduitis, chorioamnionitis, villitis and calcifications. Studies by Hirsch et al [33] and Martinot et al [35], have shown extensive placental damage in their macaque models in response to ZIKV infection showing multiple placental infarctions, chorioamnionitis and villitis. They also showed vascular pathologies including vasculitis, thrombosis and vascular collapse resulting in disturbances in transplacental oxygen transfer and decreased fetal oxygenation. Fetal hypoxia can impact fetal CNS development and may alter fetal CNS susceptibility to ZIKV. In the present study on early and mid-gestation infected pregnant baboons, placentas of both groups exhibited evidence of inflammation with increase in macrophage populations in both the maternal and fetal placental compartments compared to control. Histologically, ZIKV infected placentas exhibited several common pathological changes among the ZIKV infected dams in both early and mid-gestation groups compared to control placenta. The placentas from ZIKV infected pregnancies showed changes in maturation of villi with presence of more immature villi, disruption of the syncytiotrophoblast layer, fewer or absence of fetal capillaries in the villi, capillary thickening and thrombosis and calcium and fibrin deposits in villi and in fetal vessels. By late gestation, there is a single syncytiotrophoblast layer separating maternal and fetal compartments and any loss of integrity of this layer provides a mechanism for vertical transfer of either the virus itself or of ZIKV infected maternal immune cells. Changes in placental villi were more evident in the early-gestation than in the mid-gestation cohort. These histopathological changes have been reported in placentas of ZIKV infected women as well [52, 53]. However, we do not know what role placental inflammation and histopathological changes may have played in fetal development in our studies without detailed study of oxygenation changes on maternal and fetal side, changes in various immunological factors, placental vascular pathology and the extent to which maternal inflammation and placental pathology is transferred to the developing fetus depending on gestation age and viral presence. It is possible that fetal oxygenation was affected in the ZIKV infected dams with more severe histopathological changes to the fetal capillaries and fetal vessels that could affect the neurological development of these fetuses in absence of vertical transfer. Our previous studies in baboons also demonstrated that ZIKV infection in placenta is highly focal rather than diffused with variability amongst the various cotyledons. Therefore, it is possible that inflammation and placental function may be affected more regionally than wide spread depending on ZIKV location within the placenta and our data may not reflect the entire placenta due to sampling restriction for histology.

ZIKV infection during pregnancy causes a variety of fetal or infant neuropathology collectively known as congenital zika syndrome (CZS) described in humans and several animal models including NHPs [32, 34, 35, 38, 39]. We previously showed fetal CNS pathology after mid-gest infection in pregnant olive baboon which shed light on the timeline of vertical transfer and brain pathology after ZIKV infection and the cell types that are targeted earliest causing inflammation and damage. In this study we focused on late gestation fetal neuropathology after infection during early and mid-gestation to understand the cumulative effect of ZIKV targeting of cells at different gestation ages, chronic inflammation and the long-term effects of placental dysfunction (e.g., chronic fetal hypoxia, hypoxia/ischemia, and nutrient restriction) that are all responsible to fetal and infant CZS as reported in human population.

The clear difference between ZIKV infected fetal brains compared to control brains was observed during necropsy where structural differences in gyral and sulcal patterns were noticed in ZIKV infected brains but not control brains. Baboon cerebral development is similar to humans and in fact, of all the common NHP animal models, baboons have the largest brain with the average cerebral volume at least twice as large as that of the most commonly used NHP, the rhesus macaque, and the baboon brain was found to have the highest cerebral gyrification index (GI) and all the primary cortical structures homologues to humans [54–56]. One of the most important stages of brain development in primates is the orderly process of gyrogenesis where a smooth, lissencephalic fetal brain develops a characteristic pattern of gryri and sulcal folding [57]. Primary gyrogenesis in primates starts after completion of neuronal proliferation and migration and is accompanied by accelerated cerebral growth [54]. The features of abnormal gyri/sulci formation in the ZIKV infected fetal brains included asymmetric gyrogenesis, enlarged gyri, abnormal sulcal length and depth, absence of gyri and sulci, widening of the sulci specifically lateral and superior temporal and interhemispheric fissures (Fetus 3E and 3M) which gave these brains a misshapen and flaccid look. Abnormalities of the gyral patterns resulting in polymicrogyria, lissencephaly or pachygyria is a hallmark of ZIKV infected cases of human microcephaly [58, 59]. One of the mechanisms by which gyri/sulci form is through radial glial (RG) fibers [60]. In our previous 21 dpi acute baboon study at mid-gestation, we reported loss of RG fibers in the frontal cortex of the fetus with evidence of vertical transmission [28]. Radial glial are neural progenitor cells that are critical for cortical development in part by providing scaffolding for migrating neurons to the cortical plate from the ventricular zone via RG fibers to form the characteristic six-layered cortical structure [57]. Radial glia fibers also play an integral role in formation of sulci and gyri [61] and as such, loss of these fibers would have profound consequences on brain gyri/sulci. In our earlier study, we hypothesized that had the fetus with RG damage, and loss of RG fibers been taken to term, the loss of RG fibers and the subsequent disruption of neuronal migration during mid-gestation would have resulted in a less folded brain. Therefore, the present study showed that indeed loss of RG fibers during brain development resulted in aberrant brain folding due to gyri/sulci malformations. However, our finding of loss of certain gyri and sulci and not all supports that even in the CNS, ZIKV infection is likely restricted (similar to infecting some but not all placental cotyledons) resulting in loss of gyri and sulci in the infected areas of the cortex. Studies have shown that the size and extent of mammalian cortical folding affects cognition and sensorimotor skills and disruptions in cortical growth and folding leads to neurological disorders such as autism and schizophrenia [61–63]. Therefore, the potential for the development of long-term neurocognitive deficits in infants born with disrupted cortical growth and folding in the absence of some of the severe CZS such as microcephaly remains high. To our knowledge, this is the most extensive cortical folding disruption observed in a NHP model that emulates ZIKV induced gyral abnormalities seen in human fetus.

Histopathology using H&E staining revealed disturbances of the six-layered cortical plate (CP) in the frontal cortex in ZIKV infected fetus compared to the control brains. ZIKV infected fetal brains from early and mid-gestation groups showed focal hemorrhage in the different layers of the CP as well. Loss of pyramidal neurons in layer V of the CP was most visible in Fetus 3E (early-gest) and 3M (mid-gest) brains. Fetuses 1E, 2E (early-gest) and1M (mid-gest) had disorganization of neurons in layers IV and V where shift in pyramidal cell population from V to IV was observed. Mid-gestation Fetus 2M had disorganization of layer specific cell population where pyramidal cell population was seen in all the layers not just layers III and V. The formation of the six cortical layers (Layers I, II, III, IV, V and VI) to form the cortical plate is a sophisticated and complicated orchestration of cell migration from the germinal zone to the final destination [57]. In human fetuses, disruption of the neuronal proliferation and migration during the critical stages of neocortex development due to ZIKV infection resulted in thin cerebral cortices and in severe cases, microcephaly with partially collapsed skull [64]. In our previous acute mid-gestation baboon study, we showed dramatic loss of RG fibers in the developing fetal brain in a 21-dpi fetus which disrupted the neuronal migration to the different layers of the CP since RG fibers act as scaffolding to newly differentiated and migrating neurons to reach the many layers of the developing cortex. Therefore, this longitudinal study confirms the further disorganization and loss of cells populations in the various layers of the CP most likely through ZIKV induced loss of RG fibers during the early and mid-gestational stages by affecting neuronal proliferation, differentiation and migration.

We also observed significantly atrophied cerebellum in Fetus 3E (early-gest) and 3M (mid-gest) compared to the control brains. Cerebellum growth abnormalities such as absence, underdevelopment or atrophy are frequently seen in infants with CZS [13]. Similar to the frontal cortex, the cerebellar histological staining revealed considerably more damage to the ZIKV infected cerebellum compared to the control brains. Fetuses in both the early and mid-gestation groups showed cerebellar pathology with the most damage seen in Fetus 3E (early-gest) and 3M (mid-gest). The ZIKV infected cerebellum had significant damage to the ganglionic layer marked by distortion and loss of Purkinje cells compared to the uniform distribution of these cells in the ganglionic layer of the control cerebellum. Loss of cell volume and focal hemorrhage was also observed in the molecular and granular layers. Besides its role in motor circuits, recently several studies have focused on the non-motor circuit role of cerebellum mainly in cognitive functions such as language, behavior and emotions [65, 66]. Cerebellum’s role in several neurodegenerative (Alzheimer’s, multiple sclerosis) and neuropsychiatric diseases (Ataxia, cerebellar cognitive affective syndrome (CCAS)) have put focus on its intellectual and emotional role in the brain very much like the cerebrum due to its neural connections with important regions in the brain such as cortex, amygdala and hippocampus [65]. Children born with cerebellar hemorrhages in-utero or post-natal have language, behavioral and social deficits and neurological limitations [67]. Children born with ZIKV CZS are reported to display hyperactivity, severe irritability and self-injurious behavior which are also characteristics of behavioral disorder due to cerebellar damage [12, 67].

Similar to ZIKV induced structural damage, significant increase in Iba-1 staining, a marker for microglia, was observed in the frontal cortex and cerebellum of the ZIKV infected fetal brain than the control brains. The microglia, in situations of inflammation and injury, adopt a reactive state from their basal state by changing morphologically from a highly ramified structure to one with denser cell body with shorter thick processes and higher expression of Iba-1 than in their resting state [68]. The reactive microglia were mostly seen in the white matter of the frontal cortex and cerebellum with significantly higher Iba-1+ cells in the white matter of both early and mid-gest fetuses compared to the control brains. In the frontal cortex, the Iba-1+ cells were found to be significantly more on the early-gest brains compared to the control but not in the mid-gestation brain. This showed the non-uniform presence of inflammation in different parts of the brain due to ZIKV infection as a result of gestational age differences. Our results showed early-gestation infection had greater and long-lasting impact on brain inflammation compared to the mid-gestation infection in different areas of the brain. H&E staining of early and mid-gestation fetal frontal cortex and cerebellum had revealed vascular damage in the form of focal hemorrhage which may have induced local hypoxia/ischemia leading to inflammation. In studies of term or near-term fetal macaques, while active neuroinflammation was never reported in response to ZIKV infection, brain lesions consistent of necrosis/gliosis have been described in infants born to dams infected early in gestation [35]. While it is not certain if vertical transmission occurred in all the pregnancies in our study since no vRNA was detected in placenta or fetal tissues (either gestation age infections), we cannot ascertain if neuroinflammation observed in different brain regions was in response to direct transmission of ZIKV to the fetus or if the fetal brain inflammation could be secondary to the maternal and/or placental infection, inflammation and as such, placental insufficiency. Fetal neuroinflammation may in itself increase the risk of ZIKV infiltration to the fetal CNS through mechanisms such as leaky blood brain barrier [22, 69]. It is also possible that ZIKV may have existed in other unsampled regions of the fetal brain. Regardless of the origin of neuroinflammation, it can be postulated that the consequences of such long-lasting brain inflammation post-birth would be detrimental to brain development as witnessed in children with long-term neurological developmental defects born from mothers with ZIKV infection.

A common outcome of viral infection derived inflammation and subsequent microglial activation is increased astrocytic reactivity and gliosis. Astrogliosis have been observed in human microcephalic cases and in NHP animal models due to ZIKV infection [35, 39, 70]. Significant increase in the number of GFAP+ cells and in the intensity of the GFAP marker in these cells were seen in the frontal cortex of both early and mid-gestation fetal brains compared to the control. The reactive astrocytes undergo morphological changes in their active state by increasing in size, showing higher number of cytoplasmic processes and increasing expression of intermediate filaments especially GFAP, which is detected as increased marker intensity [71]. Astrocyte mediated inflammation due to ZIKV infection can have multiple detrimental consequences such as neuronal death and dysfunction. Vascular abnormalities such as changes in permeability is possible due to glia-mediated inflammation induced by ZIKV infection. A leaky blood brain barrier was reported in a mouse model following ZIKV infection due to changes in vessel density and diameter leading to compromised vascular integrity [22]. Blood brain barrier leakage could also lead to neuronal calcification, a key feature in microcephalic brains [59].

Oligodendrocyte development in humans starts prenatally, around mid-gestation, and continues post-birth into adulthood. Similarly, myelination, a process by which oligodendrocytes myelinate axons, takes place in late gestation and progresses prenatally over several decades based on neuronal activity and most importantly, in humans, white matter has been shown to change with experiences [72, 73]. In humans and macaques, ZIKV infection is known to cause white matter damage and reduction in myelination [20, 39, 74]. In our previous study on the acute (three week) effects of ZIKV infection at mid-gestation, we found a significant change in morphology in a subset of oligodendrocyte progenitor cell (O1+) population in the frontal cortex suggestive of oligodendrocyte damage. In the present study, we did not see changes in Olig-2+ cells (oligodendrocyte precursors) in the frontal cortex of either early or mid-gestation infected fetuses compared to the controls. It is possible that by term, numbers of the late oligodendrocyte precursors and cells expressing mature oligodendrocyte lineage markers could have recovered or been replenished [28] Therefore, other cell populations of oligodendroglial lineage at different stages of maturation need to be studied to determine detrimental changes to the mature oligodendrocyte population and subsequent myelination in late-gestation and post-birth. It is also possible that white matter injury occurred in these fetal brains, which would be difficult to identify in a fetal NHP brain without using magnetic resonance imaging (MRI). However, in the cerebellum of early-gestation infected fetuses but not in the mid-gestation, we observed a significant loss of Olig-2+ expressing cells. It is not clear why ZIKV affected Olig-2+ cell population in early-gestation infected fetal cerebellum but not mid-gestation but it is possible that ZIKV infection during early cerebellar development disrupted oligodendrocyte progenitor generation and/or differentiation into oligodendrocyte lineage. The severe damage to cerebellar structure and atrophy seen in early and mid-gestation fetuses could be in part due to damage to oligodendrocyte precursor/progenitor cell numbers and their maturation and differentiation and/or inflammation through microglial activation depending on the gestational age, when and how severe the infection occurred.

ZIKV infection decreased neural progenitor cell (NPC) population in the sub granular zone (SGZ) of the hippocampal dentate gyrus (DG) where active neurogenesis takes place throughout life unlike in the fetal cortex where neurogenesis is complete by mid-gestation in primates. We saw reduced Nestin+ cells in the DG of ZIKV infected fetal hippocampus compared to the control. Besides fewer Nestin+ cells, cells expressing Nestin appeared to be largely disorganized along the DG with shorter and distorted neuronal processes compared to the control. Loss of NPCs in the SGZ of hippocampal DG have been described in the late gestation fetal or infant macaque following ZIKV infection in early or mid-gestation [34, 39]. However, ZIKV infection did not affect actively replicating neuroprogenitor cells (Ki-67/Nestin+) or migration of neurons from DG as staining intensity of doublecortin positive (DCX+) cells were not different between ZIKV and control brains although the neuronal processes or neurites of the DCX+ cells in the ZIKV positive fetal hippocampus appeared sparse and shorter in length. It is possible that changes in progenitor population and the subsequent effect on differentiating, migrating neurons happened early on in hippocampal development during early and mid-gestation and therefore, we did not see immediate impact of ZIKV infection on hippocampal development late gestation. Inflammation was also not a factor in the hippocampus unlike in the fontal cortex and cerebellum which suggests difference in inflammation response to ZIKV infection in different regions of the brain. It is also possible that hippocampal inflammation may have been resolved by late gestation unlike other brain areas. However, any changes in NPC population as shown by our Nestin data in ZIKV fetuses could potentially have long term consequences to hippocampal neurogenesis, neuroplasticity and functional changes mainly cognition, behavior and emotional functions.

In conclusion, the longitudinal study in pregnant baboon with ZIKV infection at early and mid-gestation recapitulates the brain pathology associated with human CZS cases. While microcephaly may or may not be a possible clinical outcome in baboons due to the physiological and phylogenetic differences between humans and NHPs, our model is the first among NHPs to match many of the neuropathology such as observed in ZIKV infected cases of human microcephaly. While vertical transfers in longitudinal studies are hard to confirm, it is also the case in most human pregnancies, where most often ZIKV induced neurodevelopmental abnormalities are not obvious until birth if the fetus remains viable. Remarkably, all the fetuses sustained some CNS injury with ZIKV exposure *in utero* with those in the early gestation infection group sustaining the most on a cellular level as reported in human ZIKV cases where early-gestation infections were reported to cause the most severe CZS outcome than later gestational period. We now know from human cases that subtle fetal injuries *in utero* in the absence of microcephaly and overt CZS can still have detrimental long term cognitive, behavioral, emotional and social consequences. This study highlights the importance of studying direct or indirect inflammation in the fetal CNS, in the development of neuropathology and the mechanisms via which ZIKV induces long-term neurodevelopmental deficits. Future studies need to address viral persistence in female reproductive tract and immune privileged sites, maternal immune evasion and viral transmission dynamics in order to develop effective vaccine and therapies during pregnancy. In addition, studies of ZIKV and its ability to vertically transfer past the placental barrier to invade the fetal compartment and cause permanent damage to the brain by efficiently evading the maternal immune system, could help us better understand and prepare for future ZIKV epidemic and for potentially deadlier emerging and re-emerging teratogenic viral threats to public health.

## MATERIALS and METHODS

### Ethical Statement

All experiments utilizing baboons were performed in compliance with guidelines established by the Animal Welfare Act for housing and care of laboratory animals and conducted in accordance with and approval from the University of Oklahoma Health Sciences Center Institutional Animal Care and Use Committee (IACUC; protocol no. 101523-16-039-I). All studies with ZIKV infection were performed in Assessment and Accreditation of Laboratory Animal Care (AAALAC) International accredited ABSL2 containment facilities at the OUHSC. Baboons were fed standard monkey chow twice daily as well as receiving daily food supplements (fruits). Appropriate measures were utilized to reduce potential distress, pain and discomfort, including post-CSF collection analgesia. All animals received environmental enrichment. ZIKV infected animals were caged separately but within visual and auditory contact of other baboons to promote social behavior and alleviate stress. At the designated times post inoculation (**Fig 1**), the animals were euthanized according to the recommendations of the American Veterinary Medical Association (2013 panel on Euthanasia).

### Animals

Adult timed pregnant female olive baboons (early gest n=4 Dams 1-4E; mid-gest n=3, Dams 1-3M) were utilized for this study. All females were multiparous with history of successful prior pregnancies. All dams used in this study were determined to be seronegative for West Nile Virus prior to infection and in response to ZIKV [50].

### Virus stocks, Infection and sample collection

Animals were anaesthetized with an intramuscular dose of Ketamine (10 mg/kg) before all procedures (viral inoculation, blood, salivary and vaginal swabs and urine collection). Timed pregnant female baboons were infected subcutaneously at the mid-scapular area with a single clinically relevant dose of 10^4^ plaque forming units (pfu; 1 ml volume per dose) of the Puerto Rican ZIKV isolate (PRVABC59). The dosage used to infect the animals in our study is based on the previous works done in mosquitoes carrying WNV and DENV, where it was estimated that mosquitoes carry 1x10^4^ to 1x10^6^ plaque forming units (PFU) of the virus [75], from a study evaluating Brazilian ZIKV in a bite from *Aedes aegypti* mosquito [76] and from a study of mosquito transmission of ZIKV in rhesus monkeys [77]. The pregnant females were infected near early-gestation (between 55-58 days of gestation [dG]) and mid-gestation (between 92-97 dG; term is approx. 181 dG; the overall approach is detailed in **Fig 1**). Maternal blood samples, vaginal and salivary swabs were obtained on the day of inoculation (day 0) as well as ultrasound evaluation of fetal viability. Whole blood was collected into EDTA tubes. Saliva and vaginal samples were collected by cotton roll salivette. The sampling procedure for each dam is detailed in **Fig 1**. For early-gestation group, samples (blood, vaginal and salivary swabs) were obtained at days 0, 4, 7, 14, 21, 42, 80 and 115 post infection and amniotic fluid and maternal-fetal tissue were collected at the termination of the study (169-172 dG). For mid-gestation group, samples (blood, vaginal and salivary swabs) were collected at days 0, 4, 7, 14, 21, 42, 73 post infection and amniotic fluid and maternal-fetal tissue were collected at the termination of the study (167-169 dG). At the end of the study for each animal, dams were sedated with ketamine, all maternal samples obtained as well as ultrasound measurements, then the animal rapidly euthanized with euthasol. A C-section was quickly performed, cord blood obtained and the fetus euthanized with euthasol. Maternal and fetal tissues were rapidly collected and samples were both fixed with 4 % paraformaldehyde and frozen on dry ice (stored at -80°C) for each tissue.

Complete blood counts (CBCs): CBCs were evaluated for all females on EDTA-anticoagulated whole blood samples collected on day 0 and subsequent days-post infection as shown in the experimental timeline (Idexx ProCyte DX hematology analyzer; Idexx laboratories, ME). CBC’s included analysis for red blood cells (RBCs), hemoglobin, hematocrit and platelet count. RBC, hemoglobin and hematocrit numbers did not show any differences pre-and post ZIKV infection for any of the infected females. Platelet counts did not change in response to ZIKV infection in any dam.

### One-Step quantitative reverse transcription PCR

- Primers and probes used for qRT-PCR were designed by Lanciotti et al [78] (**Table 2**). RNA was isolated from maternal and fetal tissues (**Table 1**) using QIAamp cador pathogen mini kit (Qiagen, Valencia, CA). ZIKV RNA was quantitated by one-step quantitative real time reverse transcription PCR using QuantiTect probe RT-PCR kit (Qiagen) on an iCycler instrument (BioRad). Primers and probes were used at a concentration of 0.4 μM and 0.2 μM respectively and cycling conditions used were 50°C for 30 min, 95°C for 15 min followed by 40 cycles of 94°C for 15 s and 60°C for 1 min. Concentration of the viral RNA (copies/milliliter) was determined by interpolation onto a standard curve of six 10-fold serial dilutions (10^6^ to 10^1^ copies/ml)) of a synthetic ZIKV RNA fragment available commercially from ATCC (ATCC VR-3252SD). The cutoff for limit of detection of ZIKV RNA was 1x10^2^.

### ZIKV ELISA

ZIKA specific IgM and IgG antibody responses were assessed in the serum samples using the commercially available anti-ZIKV IgM (#ab213327, Abcam, Cambridge, MA) and IgG (#Sp856C, XpressBio, Fredrick, MD) ELISA kits. Briefly, a 1:100 for IgM and 1:50 for IgG serum dilution was performed in duplicate and added to the pre- >coated plates available in the kits. The assays were performed using the manufacturer’s instructions and the assay was read at 450 nm for IgM and 405 nm for IgG antibodies in the serum.

**Table 2.**
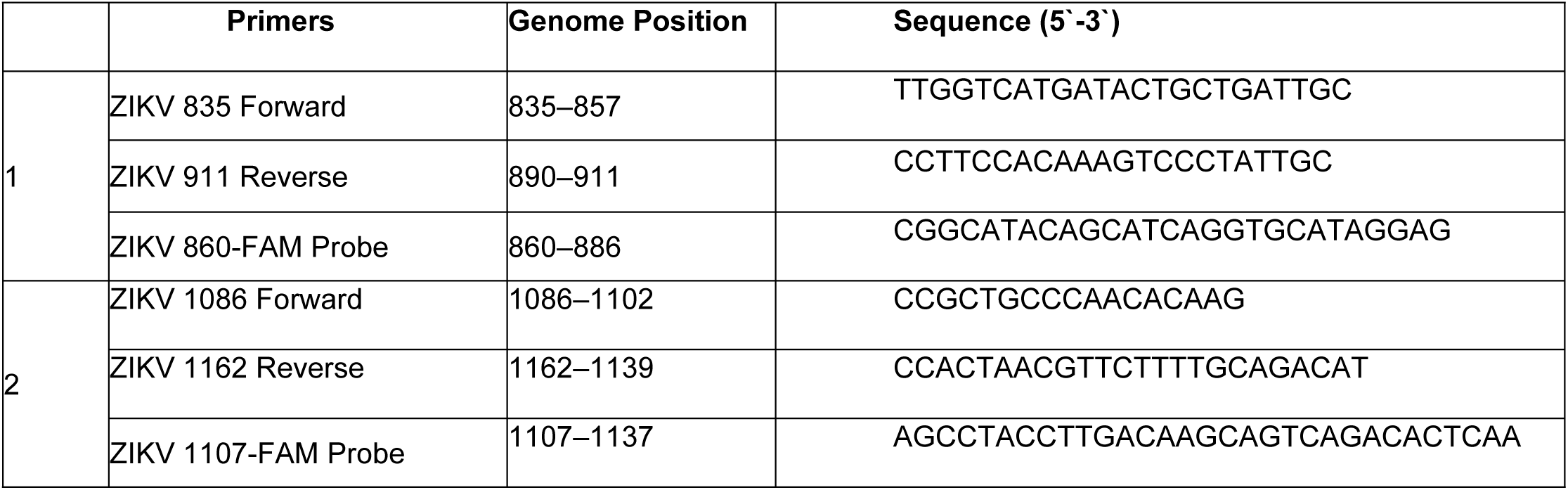
Primer/Probe sets for the detection of ZIKV by one step qRT-PCR

### Fetal CNS and placental Immunohistochemistry/Immunofluorescence

- Following removal, fetal cerebrum and cerebellum were divided mid-sagittal and frontal cortex and hippocampus dissected out of each cerebral hemisphere. one half of each tissue was rapidly frozen on dry ice and stored at -80°C and the other half fixed in 4% paraformaldehyde for 48 hours, transferred to 70% EtOH and paraffin embedded following paraffin processing and embedding protocols. The identical region of the frontal cortex was selected from each fetal brain and subjected to serial coronal sectioning (5 μm) along with the cerebellum and hippocampus. Fetuses (n=3) delivered via C-section from late gestation pregnant baboons were used as controls for this study.

Histological staining in the form of H&E stain was done on fetal frontal cortex, cerebellum, hippocampal and placental tissue sections from all ZIKV and control fetuses. Sections were visualized using a brightfield microscope (Olympus B40x) equipped with a SPOT 5 MP digital camera with SPOT 5.3 imaging software (Sterling Heights, MI).

For immunocytochemical and immunofluorescent labelling, sections every 150 microns were selected for immunolabeling for a total of 4 sections per fetus. For dual Immunofluorescence labeling (IF) on paraffin sections, after deparaffinizing and rehydrating protocol, HIER (heat-induced epitope retrieval) was performed in a Retriever 2100 with citrate-based antigen unmasking solution (#H-3300, Vector Laboratories, Burlingame, CA). After retrieval, slides were rinsed in 1XPBS and treated with 1% NaBH4 solution to reduce auto-fluorescence then blocked for two hours in blocking solution at room temperature in the humidity chamber followed by overnight incubation in primary antibodies (Iba-1 Rabbit polyclonal 1:100, NBP2-16908; Nestin; mouse monoclonal 1:100, NBP1-92717, Ki-67 Rabbit polyclonal 1:100. NB500-170 Novus Bio, Littleton, CO; NeuN, mouse monoclonal 1:200, #94403S; GFAP, Rabbit polyclonal 1:500, #12389S; DCX, Rabbit polyclonal 1:200, #4604S, Cell Signaling, Danvers, MA; Olig-2, Rabbit polyclonal 1:500, AB9610, Millipore, CA). Sections were incubated for an hour the next day at room temperature in dark in goat anti-mouse Alexa 568, goat anti-rabbit Alexa 568 and goat anti-mouse Alexa 488 1:200 (Life Technologies, Carlsbad, CA) based on the combinations of primary antibodies used for the immunostaining. After PBS washes, sections were incubated for 30 min in dark at room temperature in Prolong Gold with DAPI (Life technologies, Carlsbad, CA), cover slipped and cured for 24 h before visualizing using fluorescent microscope (Olympus BX43). Control reactions were performed where the primary antibody was omitted from the procedure. Images were captured using CellSens imaging software (Olympus Scientific Solutions, Waltham, MA).

For placenta and uterine IF, slides were baked for one hour at 56°C, deparaffinized, and HIER was performed in the Retriever 2100 with R-Universal Epitope Recovery Buffer (62719-10 lot 180314). After retrieval, slides were blocked in 5% normal donkey serum for 1 hour, then primary antibodies in 0.5% normal serum were added and incubated overnight, humidified, at 4°C [MAC-387, macrophage antibody (Abcam, MA); Pan anti-flavivirus; (Millipore, CA)]. The next morning slides were removed from 4°C and allowed to equilibrate to RT, covered, on the benchtop for 1 hour. Slides were rinsed 4 x 5 minutes with PBS, then secondary antibodies were added and incubated 1 hour, covered, at RT. Donkey anti-mouse IgG F(ab’)2 AlexaFluor 594 (Jackson Immunolabs) was used as secondary antibody. Slides were rinsed in PBS, counterstained 5 minutes with DAPI in PBS and cover slipped using Shur/Mount. Cover glass were sealed with nail polish and slides were stored at 4°C and visualized using a fluorescent microscope (Olympus BX43). Images were captured using CellSens imaging software (Olympus).

### Image analysis

Image analysis was performed using NIH ImageJ FIJI (Wayne Rasband, NIH). Original IF images were split into separate channels (Red, green, blue) using identical parameters for each section (minimum, maximum, brightness, contrast). Using LUT, immunostained cells were given a different color to separate from DAPI channel. After the images were restacked to final composite image, using cell counter plugin, cells were counted manually, then total cells for each section were counted. For Nestin positive cells, NeuronJ plugin in FIJI was used to trace the neurites and the sum of length of all tracings for each section was measured. For DCX+ cell analysis, images were converted to Binary and then skeletonized. 5 fields along the DG were then selected and mean gray value (MGV) of these regions of interest (ROI) and the background was measured. Specific MGV was calculated as the difference between the background MGV and that of each ROI on DG. MGV is a measure of staining intensity in an area, in this case, directly corresponding to DCX+ cell staining along the DG of the hippocampus.

## ACKNOWLEDGEMENTS

This research was supported by the NIH NINDS R21NS1037720-01, The Oklahoma Baboon Breeding Resource; The Office of the Vice President for Research, University of Oklahoma Health Sciences Center; Oklahoma Medical Research Foundation (OMRF) Imaging Core Lab for help with paraffin processing and histology.

